# Nickel-Sulfonate Mode of Substrate Binding for Forward and Reverse Reactions of Methyl-SCoM Reductase Suggest a Radical Mechanism Involving Long Range Electron Transfer

**DOI:** 10.1101/2021.01.25.428124

**Authors:** Anjali Patwardhan, Ritimukta Sarangi, Bojana Ginovska, Simone Raugei, Stephen W. Ragsdale

## Abstract

Methyl-coenzyme M reductase (MCR) catalyzes both synthesis and anaerobic oxidation of methane (AOM). Its catalytic site contains Ni at the core of Cofactor F_430_. The Ni ion, in its low-valent Ni(I) state lights the fuse leading to homolysis of the C-S bond of methyl-coenzyme M (methyl-SCoM) to generate a methyl radical, which abstracts a hydrogen atom from Coenzyme B (HSCoB) to generate methane and the mixed disulfide CoMSSCoB. Direct reversal of this reaction activates methane to initiate anaerobic methane oxidation. Based on crystal structures, which reveal a Ni-thiol interaction between Ni(II)-MCR and inhibitor CoMSH, a Ni(I)-thioether complex with substrate methyl-SCoM has been transposed to canonical MCR mechanisms. Similarly, a Ni(I)-disulfide with CoMSSCoB is proposed for the reverse reaction. However, this Ni(I)-sulfur interaction poses a conundrum for the proposed hydrogen atom abstraction reaction because the >6 Å distance between the thiol group of SCoB and the thiol of SCoM observed in the structures appears too long for such a reaction. Spectroscopic, kinetic, structural and computational studies described here establish that both methyl-SCoM and CoMSSCoB bind to the active Ni(I) state of MCR through their sulfonate groups, forming a hexacoordinate Ni(I)-N/O complex, not Ni(I)-S. These studies rule out direct Ni(I)-sulfur interactions in both substrate-bound states. As a solution to the mechanistic conundrum, we propose that both forward and reverse MCR reactions emanate through long-range electron transfer from Ni(I)-sulfonate complexes with methyl-SCoM or CoMSSCoB, respectively.

---

Methyl-coenzyme M reductase (MCR), one of the few Ni proteins in nature, catalyzes the reaction of methyl-coenzyme M (CH_3_-SCoM) with coenzyme B (HSCoB) in methanogenic archaea to form methane and the heterodisulfide, CoMSSCoB (Eq. 1) (1). MCR also catalyzes the reverse reaction in consortia of anaerobic methane oxidizing archaea (ANME) with sulfate, nitrate or Fe(III) reducing bacteria (2). This manuscript describes the relative orientations of the substrates in the forward and reverse reactions of MCR. Spectroscopic, structural and computational studies identify an unexpected and symmetric complex between active Ni(I) enzyme and the sulfonate group of the substrates for the forward and reverse reactions of Eq. 1. These results suggest the need to reassess how catalysis is triggered in the MCR mechanism.

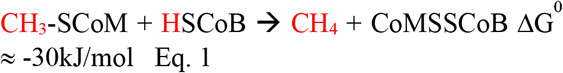

Understanding the biosynthesis of methane is important from basic energy, economic, and environmental perspectives. Methane accounts for 22% of U.S. energy consumption (3,4), with half of homes using natural gas as their heating fuel. Methane is the simplest organic compound, but it has the highest energy content of any carbon-based fuel. About 90-95% of all methane on earth is produced biogenically (5). Methanogens, responsible for enzymatic synthesis of one billion tons of methane per year (6), are its major source, the balance generated by metabolism of methyl-mercury (7) and methylphosphonate (8). Much of the methane formed by methanogens is captured and used as an energy source by aerobic and anaerobic methanotrophic (ANME) microbes (9,10). However, the increased mining of methane and industrial farming of cattle and other ruminants since the industrial revolution has created a mismatch between the sources and sinks of methane, causing its atmospheric levels to double over the past two centuries (11). This is an environmental concern related to global warming because methane is 21 times more effective at trapping heat than the other major greenhouse gas, CO_2_ (12).

The basic chemistry and biology of alkane activation and formation by MCR is also intriguing because of methane’s strong sp^3^ C–H bonds, its low solubility in both polar and nonpolar solvents, and very high ionization energy. The selective activation and functionalization of the C-H bond of methane was dubbed one of the holy grail reactions in chemistry (13). MCR accomplishes this reaction, as well as methane synthesis, using a reduced Nihydrocorphin cofactor F_430_ (14-17) related to porphyrin, chlorophyll, and vitamin B_12_, along with second sphere amino acid residues, as catalysts. Computational (18) and experimental (19) results indicate that the methane formation and oxidation by MCR occur through a mechanism involving methyl, thiyl and disulfide anion radicals.

A variety of crystal structures of MCR in the presence (and absence) of substrates have been published (20-24). It is a multimeric enzyme composed of three subunits MCRA (α), MCRB (β) and MCRG (γ) in an α_2_β_2_γ_2_ conformation with a reduced hydrocorphin Ni cofactor F_430_. buried at the end of a 30 Å long substrate channel. The F_430_ is anchored via a combination of electrostatic and H-bond interactions between the carboxylate side chains and the protein backbone (**Figure 1)**. A weak distal Ni-O bond (O-α’Gln) is present in all forms of the Ni(II) enzyme as well as the Ni(I) states, MCRred1, MCRred1-silent and MCR-Me (20,25). The coordination environment of the different states of MCR are shown in **Figure 2**. The Gln residue moves in (2.1 Å, 6 coordinate) or away (2.3 Å 5 coordinate) based on the presence or absence of a proximal axial ligand to the Ni-center and may play a role in fine tuning the redox potential of the Ni-center and/or provide stability during catalysis (20). Two moles of F_430_ bind the hexameric protein at identical but distinct substrate binding sites separated by 50 Å. The catalytic Ni site is accessible only through this channel and only to small molecules up to a diameter of 6 Å (20).

**Figure 1.**
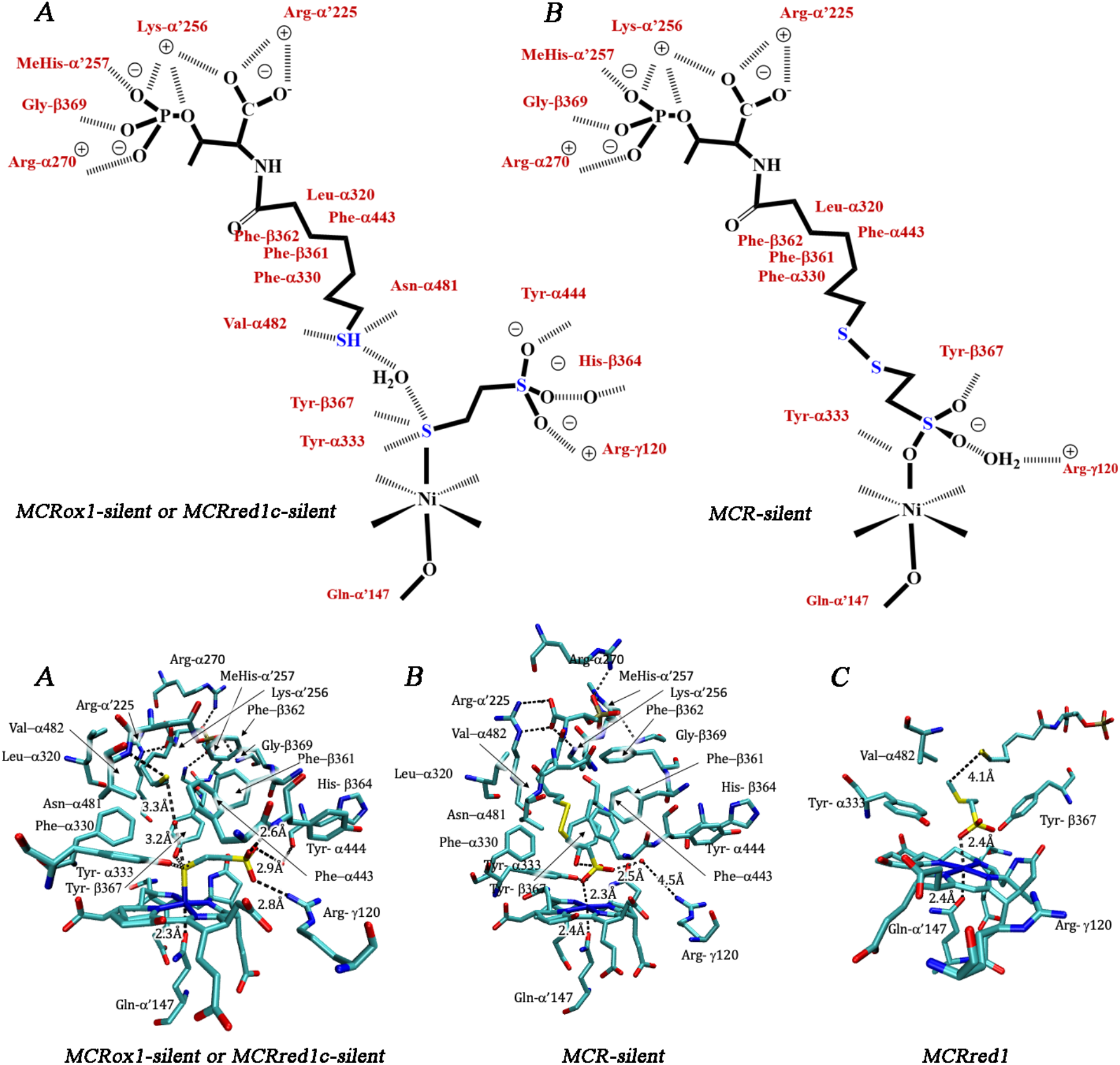
Representative interactions in the active site of MCR. Top. A: MCRox1-silent or MCRred1c-silent. B; MCR-silent. CoB part of the molecule in both A and B are stabilized at the top of channel by electrostatic interactions with phosphate and carboxylate residues. The methylene [(CH_2_)_6_-CO} section of CoB interacts with hydrophobic residues of the protein. H-bonds to the S-of CoMSH in the reduced form (A) are replaced by H-bonds to the sulfonate –O-of the CoM moiety in CoMSSCoB (B). In A, Coenzyme M is anchored to the protein by the negatively charged sulfonate group, forming a salt bridge to the guanidinium group of Arg-γ120, a hydrogen bond to Tyr-α444, and a hydrogen bond to a water molecule connected to the peptide oxygen of His-β364. In B, one oxygen atom of the sulfonate is axially coordinated with the nickel and contacts the hydroxyl group of Tyr-α333. The second oxygen atom is hydrogen-bonded to the lactam ring of F_430_ (not shown) and to the hydroxyl group of Tyr-β367 and the third to a water molecule located at the former binding site of the sulfonate in A (20). Structure details obtained from pdb1mro. Bottom. A. MCRox1-silent or MCRred1c-silent. B. MCR-silent. C. MCRred1 with methyl-SCoM and CoBSH modeled into the active site showing the predicted distances between the S-methyl carbon and the HSCoB hydrogen atom to undergo abstraction. In figure C, F430 and all residues except for methyl-SCoM and HS-CH_2_-CH_2_-tail of HSCoB are from pdb1hbm. The calculated QM model (Fig 10(3)) was overlapped with pdb1hm based on the subset of atoms fixed in the crystal structure position during the optimization, and all atoms except for the ones representing the substrates were removed.

**Figure 2:**
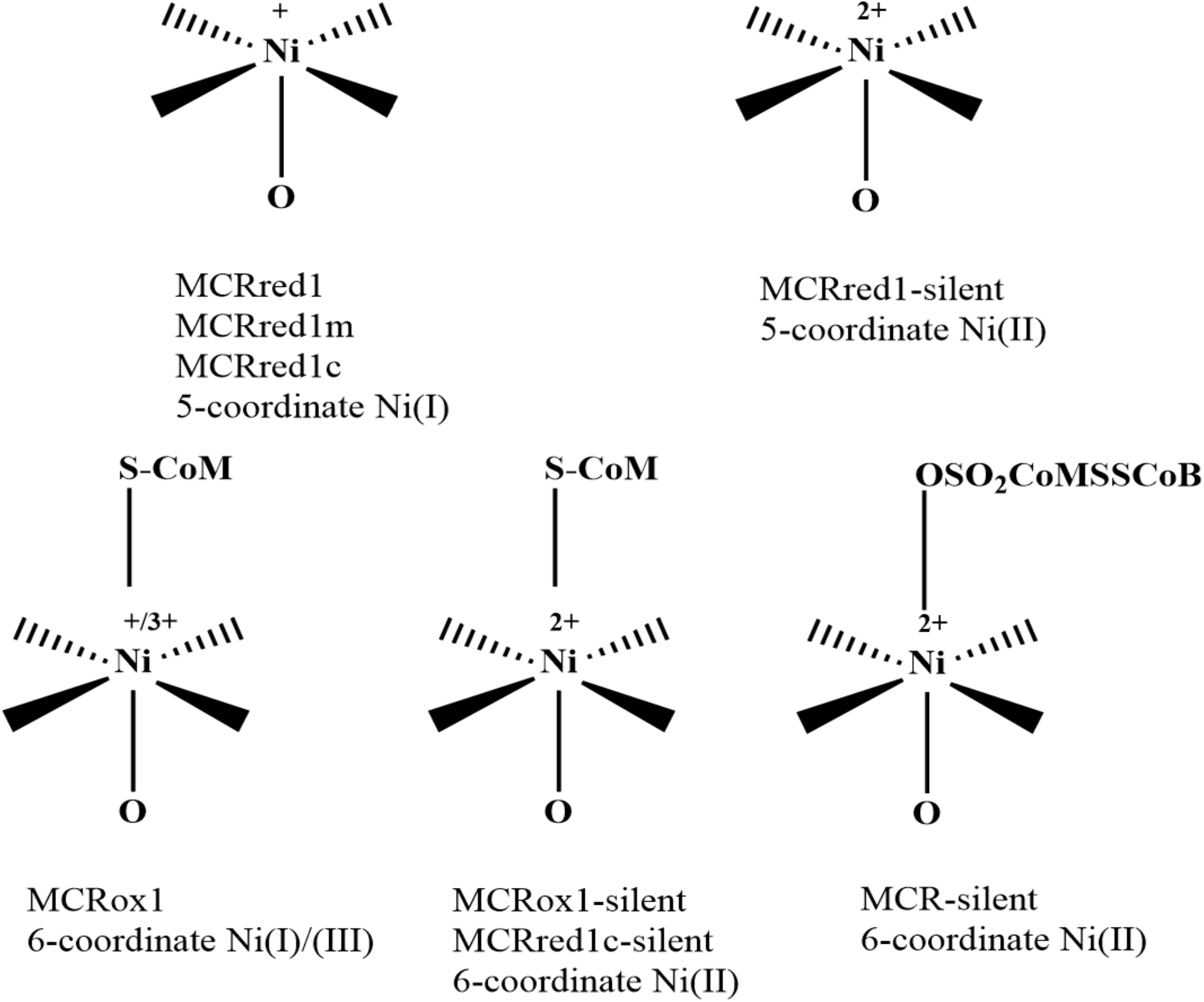
Various states of MCR, with their observed oxidation states and coordination geometries. MCRred1m: MCRred1 with methyl-SCoM, MCRred1c: MCRred1 with coenzyme M, MCRox1: MCR in the ready state with Ni in the 1+/3+ oxidation, MCRred1-silent, MCRred1m-silent, MCRred1c-silent, MCRox1-silent are the corresponding Ni^2+^ states of the Ni^1+^ states, MCR-silent is the heterodisulfide (CoMSSCoB) bound state observed by X-ray crystallography.

Regarding binding of CoM, the crystal structures of the Ni(II)-MCRred1-silent and MCRox1-silent states show its thiolate anchored by binding axially to the Ni(II) on one side and its negatively charged sulfonate group forming a salt bridge to the guanidinium group of Arg-γ120 (20,21) (Figure 1). XAS results similarly indicate S coordination to the Ni in the MCRox1 and MCRox1-silent states, while the MCR-silent state (with CoMSSCoB bound) fits best with 4N ligands and 2N/O ligands in the axial position to give a hexacoordinate Ni-center (25,26) (Fig. 2). Ni K pre-edge and EXAFS data and time-dependent DFT (TD-DFT) calculations also reveal the axial Ni-S bond from the thiolate of CoMSH in the Ni(II) MCRred1-silent and MCRox1-silent states (27). Additionally, this Ni-thiolate appears to interact with a water molecule that bridges CoMSH and HSCoB and forms hydrogen bonds to the hydroxyl groups of Tyr-α333 and Tyr-β367 (21) (Figure 1); however, Grabarse et. al speculate that due to the extremely hydrophobic nature of the pocket there cannot be a solvent molecule trapped between the CoMS-Ni(II) and HSCoB in the MCRox1-silent structure but rather a methyl radical generated during X-ray exposure of the crystal (21).

HSCoB hangs down the substrate binding channel tethered by electrostatic interactions between its negatively charged threonine phosphate moiety and five positively charged amino acids. Its heptanoyl [(CH_2_)_6_-CO-] arm stretches over 16 Å in Van der Waals contact with several hydrophobic residues, namely Phe-α330, Tyr-α333, Phe-α443, Phe-β361, and Tyr-β367 positioned along the substrate binding pocket (Figure 1), At its end, the HSCoB thiol group is 8.8 Å from the Ni-center and interacts with the side chain nitrogen of Asn-α481, the main chain peptide nitrogen of Val-α482, and a bridging water molecule. Asn-α481 is within hydrogen bond distance of the sulfur in the post translationally modified thioglycine (TGly)-α445 (20).

Crystal structures show that various HSCoB_5-9_ analogs are anchored at an annulus of charged residues at the top of the substrate channel and adopt similar positions as they thread into the channel toward the Ni-cofactor (24). The thiol group of the slow substrate HS-CoB_6_ situates at the same position as that of the native substrate, i.e. 8.8 Å from the Ni, leaving a 6.4 Å gap between the thiol group of HSCoB and the Ni-bound S of CoM. Since a methyl group of CH_3_-SCoM cannot bridge this large gap (28), a conformational change was proposed to allow HSCoB to penetrate deeper in the substrate channel towards the Ni ion. The HSCoB_5_ thiol group rests 9.3 Å from the Ni as would be expected, while the SH of CoB_8_SH/CoB_9_S are located 5.9 – 6.2 Å from the Ni of F_430_. The crystal structures of MCR with the CoBxSH analogs did not reveal any of the conformational changes proposed to occur during catalysis, suggesting a more important role for methyl-SCoM, CoMSH and the oxidation states of the Ni-F_430_ in the catalytic cycle (24) (Figure 2). Furthermore, the structure of the MCR-CoMSSCoB (MCR-silent) reveals the HSCoB portion in virtually the same place as in MCRox1-silent (29,30). Thus, the large distance in these structures (24,29,30) between the HSCoB hydrogen atom to be abstracted and the methyl group, which is proposed to undergo homolytic fission promoted by a Ni(I)-SCoM interaction, poses a conundrum for all published MCR catalytic mechanisms, highlighted in **Figure 3a**.

**Figure 3.**
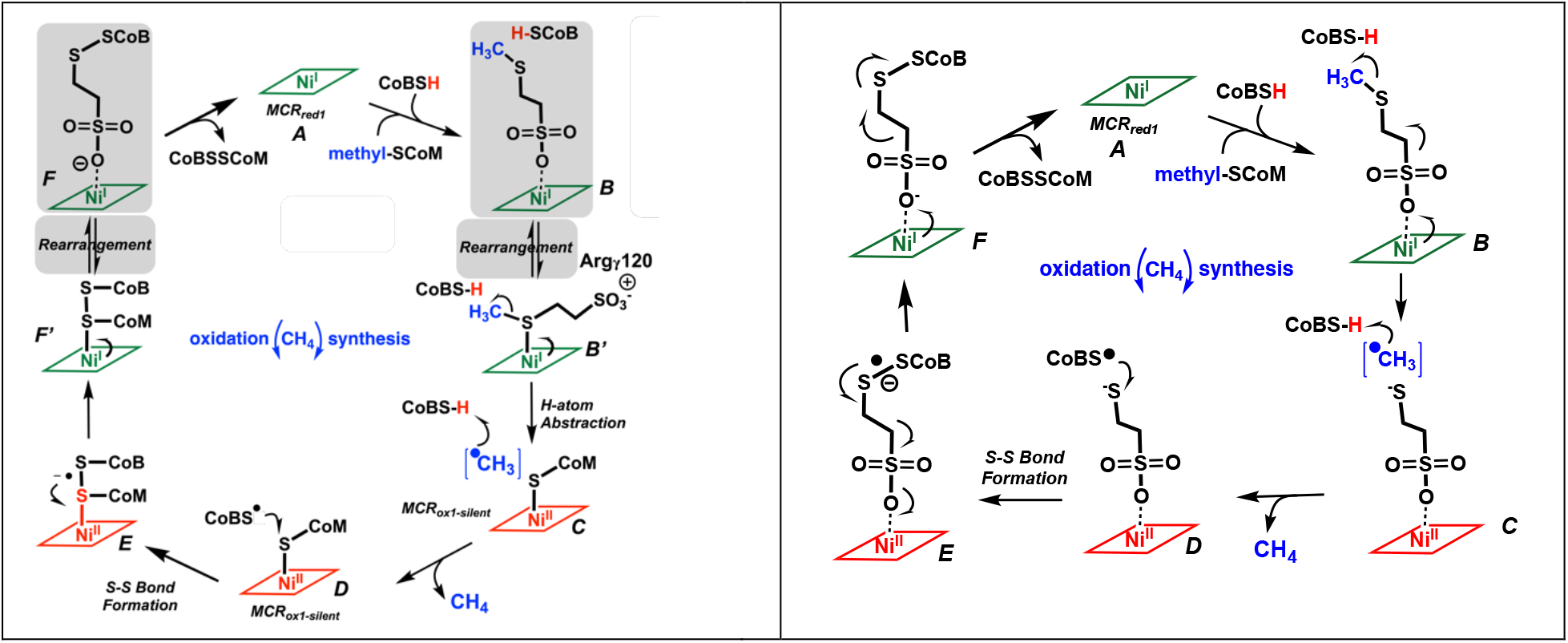
Contrasting mechanisms of activation of methane synthesis. (Left) Canonical mechanism through Ni(I)-S species with direct Ni→sulfur ET. The shaded portion includes the sulfonate-bound substrates, which could rearrange to form Ni-thiol (B’ or F’) states proposed to be productive state in the canonical mechanism. (Right) Proposed mechanism of CH_4_ synthesis involving Ni-sulfonate species (B) as productive complexes and long-distance ET. See text for details.

One significant issue is that all published crystal structures are of inactive Ni(II) or a methyl-Ni(III) state—none are available for the active Ni(I) state. In our studies of the binding of HSCoB analogs (24), although the crystallizations were set up with the MCR solution predominantly in the Ni(I)-MCRred1 state, by the time X-ray diffraction data were collected they had undergone oxidation to the Ni(II)-MCRred1-silent state, as assessed by single crystal UV-visible microspectrophotometry; furthermore, following data collection there was no evidence for photoreduction of the Ni(II) back to Ni(I) in any of the crystals. Attempts to photoreduce the crystals using different wavelengths and temperatures were also unsuccessful.

Thus, spectroscopic studies are crucial to obtaining an accurate description of the coordination chemistry of the Ni(I) state. In this respect, X-ray absorption spectroscopy (XAS) is highly significant because it provides extremely precise measurement of the Ni oxidation state and metal-ligand bond distances as well as discrimination between thiolate and nitrogen or oxygen ligation. Furthermore, these experiments can be carried out under strictly anaerobic conditions. Similarly, electron paramagnetic resonance (EPR)-based studies provide an independent view of the d^9^ Ni(I) state. For example, on the basis of EPR, electron nuclear double resonance (ENDOR) and quantum mechanical calculations, Hinderberger et.al, proposed that the thioether-S of methyl-SCoM and one of the –H atoms of the methyl group are between 3.45-3.75Å and 5.35 – 5.65Å respectively from the Ni-center (31). This binding mode does not bring the thioether–S within bonding distance of the Ni. In this contribution, we add near infrared (NIR) spectroscopy to characterize the active Ni(I) state of MCR. The spectroscopic, kinetic, structural and computational studies described here establish that methyl-SCoM binds to the active Ni(I) state of MCR through its sulfonate, forming a hexacoordinate Ni(I) complex. This is unusual— most Ni(I) complexes are four- or five-coordinate (32); we are aware of only one other structurally characterized six-coordinate model Ni(I) complex (33). Here we also suggest a solution to the apparent conundrum of how the methyl radical can attack the H of HSCoB. As shown in Fig. 3b, we propose a revised mechanism for MCR catalysis in which methyl-SCoM binds to Ni(I) through its sulfonate group. We suggest that this is the state from which catalysis ensues.

## RESULTS

### Changes in near infrared bands on addition of substrates: Comparison among the various substrates

In the UV-visible region, Ni(I) has a characteristic absorbance at 385 nm which changes to 420/445(sh) nm on oxidation to Ni(II) or Ni(III). Ni(I)-MCRred1 also exhibits characteristic d-d transition in the near infrared (NIR) region of the electromagnetic spectrum that are absent in Ni(II) states like MCRred1-silent. Addition of substrates, CoMSSCoB and methyl-SCoM to 50 µM Ni(I)-MCRred1 (between 65-70 % Ni(I) for various purifications) elicits significant changes in these NIR bands over a timeframe that the 385 nm UV-visible band associated with Ni(I) does not change. Thus, we attribute these changes to shifts in the d-d orbitals of Ni(I) that can be monitored to elucidate the nature of substrate binding.

Addition of HSCoB alone does not elicit NIR changes. However, titration of Ni(I)-MCRred1 (50 µM) with the first substrate in the reverse reaction, CoMSSCoB or CoMSSCoB_6_ (0-500 µM), shifts the broad 700 nm peak to ∼750 nm (**Figure 4A**). The breadth of these d-d transitions makes it difficult to precisely monitor the wavelength maxima due to overlap. We employed difference spectra to more accurately quantify these changes at the exact wavelengths. Difference spectra (**Figure 4B**) show that the decrease in absorbance at 700 nm is associated with an increase at 768 nm and 850 nm for both substrates. The absorbance at 385 nm characteristic of Ni(I) in MCRred1 remains unchanged as the NIR spectra shift demonstrating that the decrease in absorbance at 700 nm is due to binding and not due to oxidation of Ni(I) (**Figure 4C**) associated with catalysis. Oxidation of Ni(I) to Ni(II) occurs over a longer time frame associated with decay of the NIR bands as the 385 nm band shifts to 420 nm (**Figure S5**) Thus, these NIR changes can confidently be assigned to d-d changes associated with substrate binding alone.

**Figure 4:**
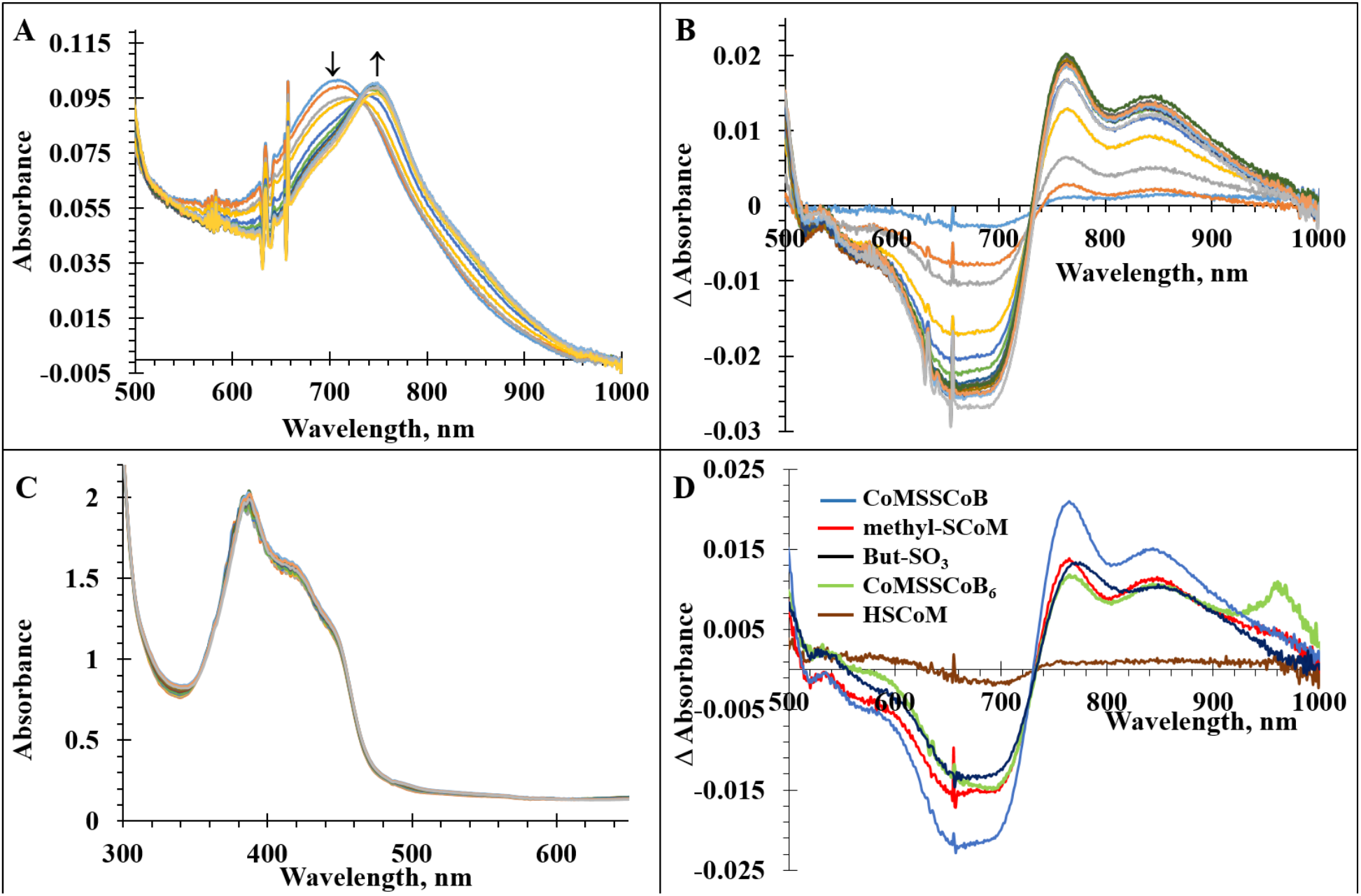
Binding substrates monitored by NIR spectroscopy. A: Red shift in λmax from 700 nm to 750 nm of Ni(I)-MCRred1(50 µM) with increasing concentration of CoMSSCoB (0-700 µM). B: Difference spectra of the spectra in panel A, highlighting the buried peaks at 768 nm and 850 nm on CoMSSCoB addition. C: The absorbance at 385 nm (Ni(I)) and 420 nm Ni(II)) remains unchanged during the addition of the substrate suggesting no redox reaction at the Ni(I) center. D: Comparative difference spectra for the addition of 500 µM of CoMSH (green), Methyl-SCoM (red), CoMSSCoB (Blue) and CoMSSCoB_6_ (yellow) and But-SO_3_ (black) each to MCRred1 (50 µM).

Interestingly, addition of methyl-SCoM to MCRred1 gives rise to the same NIR spectral changes with the absorbance at 385 nm remaining unchanged. These changes in d-d transitions suggest that CoMSSCoB, CoMSSCoB_6_, and methyl-SCoM all bind in a similar fashion within the enzyme pocket.

On the other hand, addition of CoMSH does not result in any NIR spectral shifts. Ni(I)-MCRred1 is purified in the presence of 10 mM CoMSH to stabilize the active enzyme. These preparations result in a mixture of Ni(I) and Ni(II), which can be precisely quantified by the relative absorbances of these states. The Ni(II)-MCRred1 state contains tightly bound Ni(II)-SCoM that is difficult to remove by dialysis, leading to a quantifiable proportion of the enzyme containing a Ni(II)-thiolate with the sulfonate bound to Arg-γ120 as observed in various crystal structures (described above). Previous XAS studies have shown Ni(I) in MCRred1 to be 5-coordinate in the presence of CoMSH (25). Thus, the lack of NIR changes upon addition of CoMSH results from the stability of five-coordinate Ni(I)-MCRred1 or an alternate binding conformation in the substrate pocket.

Our interpretation of these substrate-induced spectral changes is that addition of HSCoB alone or CoMSH does not result in any change in the Ni(I) d-d bands, suggesting that these substrates do not alter the five-coordinate Ni(I) state. On the other hand, addition of methyl-SCoM or either of the heterodisulfide substrates results in formation of a similar six-coordinate Ni(I) state. These conclusions are validated by XAS and computational results, described below.

Quantum chemical truncated models (**Figure 5**) of the MCR active site containing Ni(I) binding either CoMSSCoB (model 1) or methyl-SCoM (the latter with or without CoBSH, models 2 and 3, respectively) were investigated to elucidate the Ni-sulfonate interaction and its effect on the NIR spectra. Based on the p*K*_a_ measurements reported below, the sulfonate groups of methyl-SCoM and CoMSSCoB were modeled as deprotonated. Because all models truncated the SHCoB at the second carbon, the protonation of the phosphate group does not affect these calculations. The geometries of the optimized structural models are reported in Figure 5 along with relevant geometrical parameters. Additional geometrical parameters are summarized and compared with those from the 1.8 Å resolution crystal structure of the MCR-product complex (pdb entry: 1HBM) in **Table 1**.

**Table 1.** Characteristic bond distances for the crystal structure and the three models in Figure 5 calculated using gas phase DFT approach

| Bond distance (Å) | 1hbm | CoMSSCoB+CH<sub>4</sub> | CoBSH-methyl-SCoM | methyl-SCoM |
| --- | --- | --- | --- | --- |
| Ni-O(Gln147) | 2.35 | 2.03 | 2.03 | 2.04 |
| Ni-O(SO<sub>3</sub><sup>-</sup>) | 2.29 | 2.41 | 2.41 | 2.71 |
| Ni-S(SO<sub>3</sub><sup>-</sup>) | 3.29 | 3.42 | 3.45 | 3.62 |
| Ni-S<sub>m</sub> | 6.71 | 7.04 | 6.09 | 6.98 |
| Ni-S<sub>c</sub> | 8.46 | 8.47 | 8.48 | n/a |
| C(Me)-S<sub>c</sub> | n/a | 4.41 | 3.72 | n/a |

\*S<sub>m</sub>, thioether sulfur from methyl-SCoM; S<sub>c</sub>, thiol sulfur from CoM. See Fig. 5.

**Figure 5.**
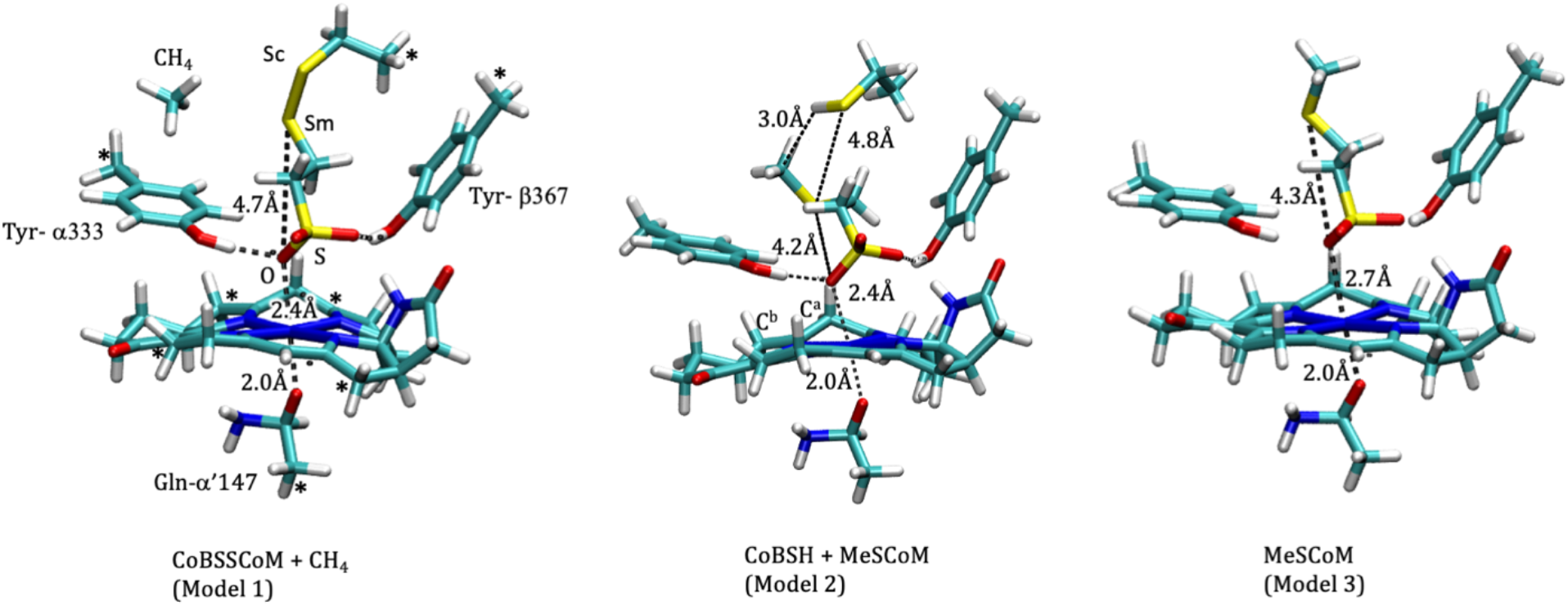
Structures optimized using DFT. 1) Ni(I) binding CoMS-SCoB, where CoB is truncated after the second carbon from the S_b_, 2) Ni(I) binding methyl-SCoM and 3) Ni(I) binding CoBSH and methyl-SCoM. All models were build starting from crystal structure 1HBM. Truncation points from the enzyme are annotated with asterisks and were fixed in place during the optimization.

All models show stable complexes of Ni with the SO_3_^-^ group and Gln147 residue via oxygen, as well as the interaction of the substrate with the OH groups of the Tyr residues,

The spectra of the structural models (**Figure 7**) for Ni coordination with SO_3_^-^ of methyl-SCoM considered in the present work were simulated in a time-dependent-DFT (TDDFT) framework and compared with the spectra of previous structural models for a Ni-S_m_ coordination (**Figure 6**, models A-D). Figure 6 shows that the adopted computational setup captures well the major spectral features and, importantly, qualitatively captures the shifts in the various structures. For structures A and B, which are Ni(I), the characteristic peak is calculated to be around ∼340 nm. As the S-S bond is broken, the S_m_-Ni bond is formed (structures C and D) and Ni center changes from Ni(I) to Ni(II) as the peak shifts towards ∼460 nm, in agreement with the experimentally observed shifts. In particular, the d-d transitions for structures A and B are captured at around 650 nm (**Figure 6b**). For the structures with Ni-O coordination (Figure 7 and **8**), the d-d transitions in the NIR spectrum are slightly shifted to ∼600 nm, with some low intensity d-d transitions found beyond 780 nm. These calculations suggest that the spectral features observed experimentally can be assigned to the structures exhibiting Ni-SO_3_^-^ interactions.

**Fig. 6.**
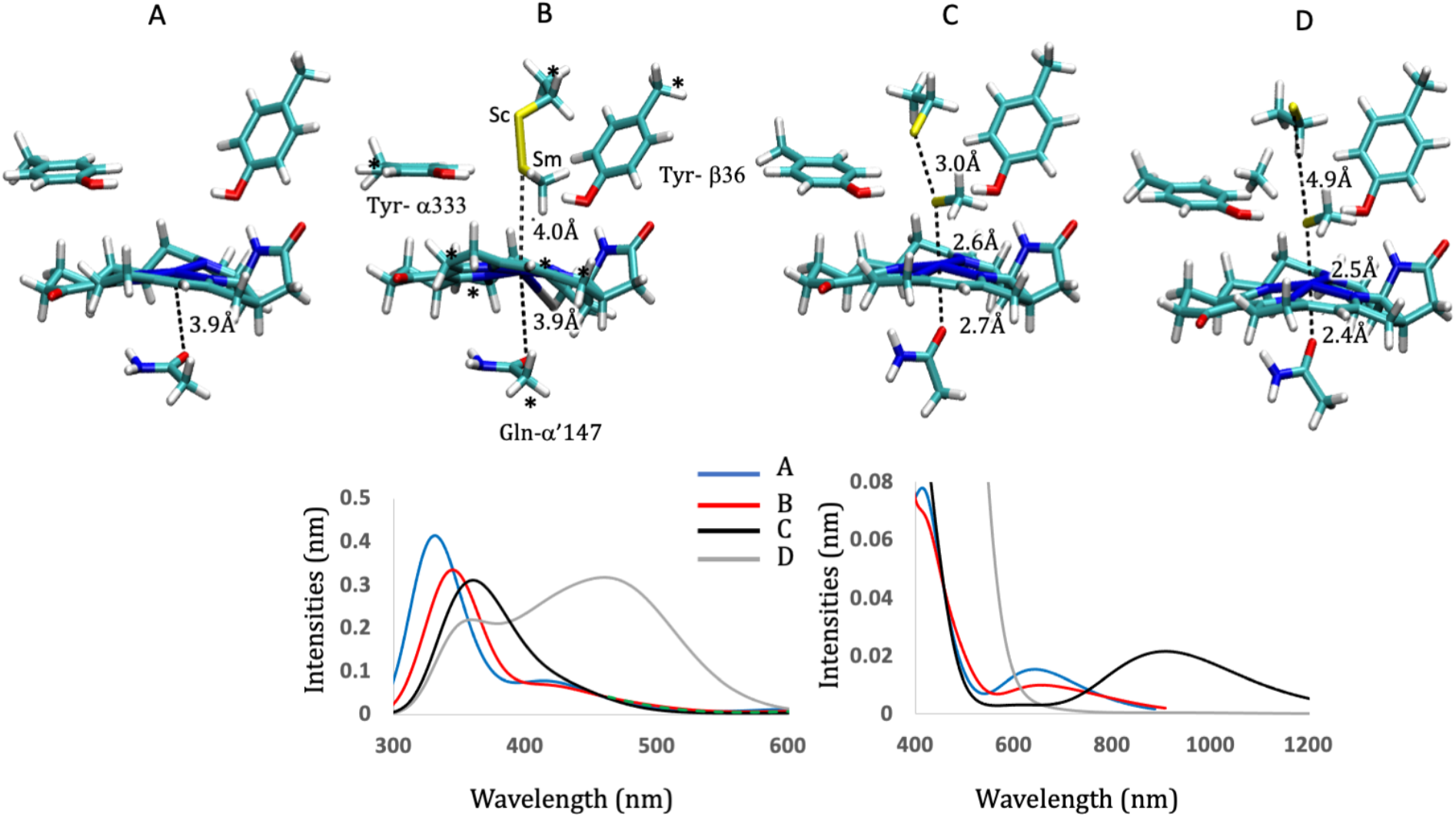
Computational models of the canonical mechanism involving Ni-S_m_ coordination and TDDFT calculations of MCR UV-vis and NIR spectra. (B) and bond formation (C and D). All models were previously reported in Ref (38). Structures A and B are Ni(I) state, and structures C and D are Ni(II). (a) Major features of the spectra as captured by the TDDFT calculations for all models shown on the bottom panels and are able to qualitatively capture the shifts in the various structures (bottom, left panels). (b) shows that d-d transitions are captured for structures A (MCRred1) and B (MCRred1 + CoMSSCoB) at around 650 nm (bottom, right panel).

**Fig. 7.**
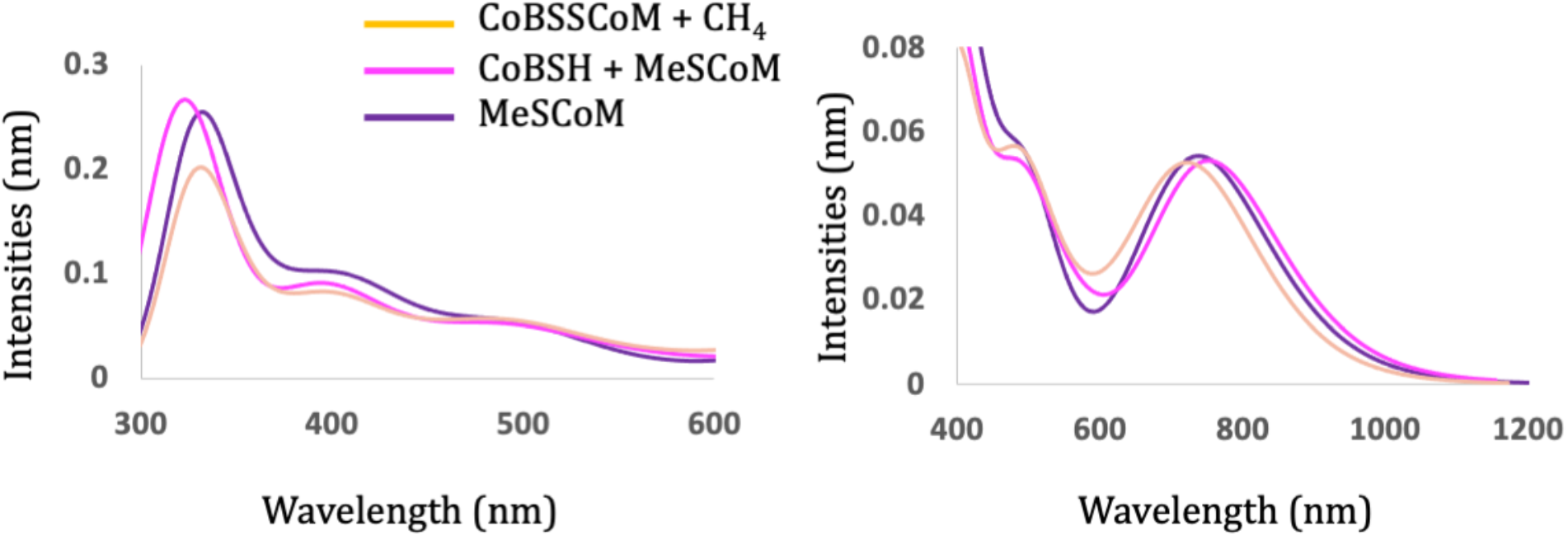
TDDFT calculations of MCR UV-vis and NIR spectra. (a) Major features of the spectra as captured by the TDDFT calculations for the models shown in Fig. 5, are able to qualitatively capture the shifts in the various structures (left panels). (b) shows that d-d transitions are captured for all three structures ∼600 nm, and extend beyond 800 in the NIR region (right panel). The more prominent peak at ∼740 nm captures ***π* to *π*\*** on F_430._

**Fig. 8.**
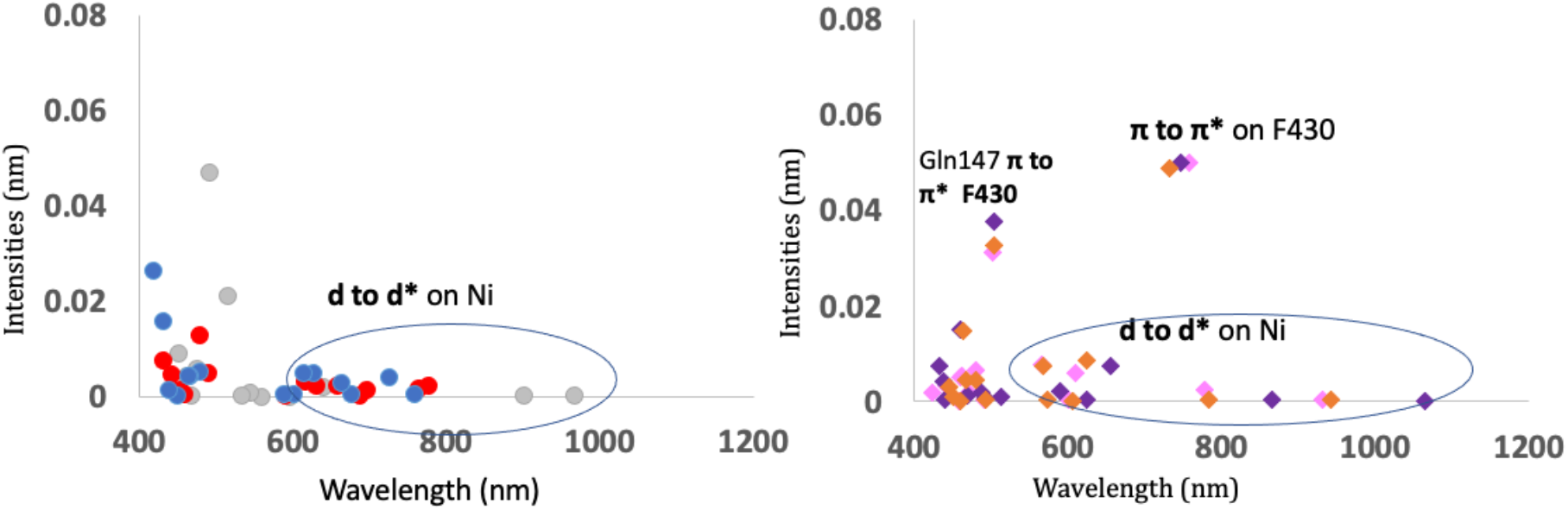
Excitation from TDDFT used in reconstructing the spectra. The calculated transitions used to reproduce the spectra in the d-d region for all models shown in Fig. 5 (right panel) and Fig. 6 (left panel).

### Binding constants for methyl-SCoM, CoMSSCoB and CoMSSCoB6

A plot of the NIR changes shows that the decrease in absorbance at 700 nm matches the concomitant increase at 768 nm and 850 nm for CoMSSCoB, CoMSSCoB_6_, and methyl-SCoM (**Figure 9A**) The data are fit to a one-site binding isotherm, where fraction of bound enzyme is plotted versus the concentration of substrate. For the reverse reaction substrates, the K_d_ values are 57.4 ± 5.4 µM for CoMSSCoB and 105.3 ± 10.9 µM for CoMSSCoB_6_ (**Figure 9B**). The K_d_ for methyl-SCoM is 27.5 ± 7.3 µM (**Figure 9C**), which is similar to the value (13 ± 4 µM) obtained by fluorescence spectroscopy (34). The tighter binding for methyl-SCoM is as expected, since MCR relies on the strict binding order of substrates to initiate catalysis. The K_d_ values for the heterodisulfide substrates are similar to that for HSCoB (90 ± 22 µM) as measured by fluorimetry (34), presumably dictated by the strong electrostatic interactions at the top of the channel between the positively charged protein residues (Figure. 1) and the carboxylate and phosphate groups of the CoB moiety.

**Figure 9:**
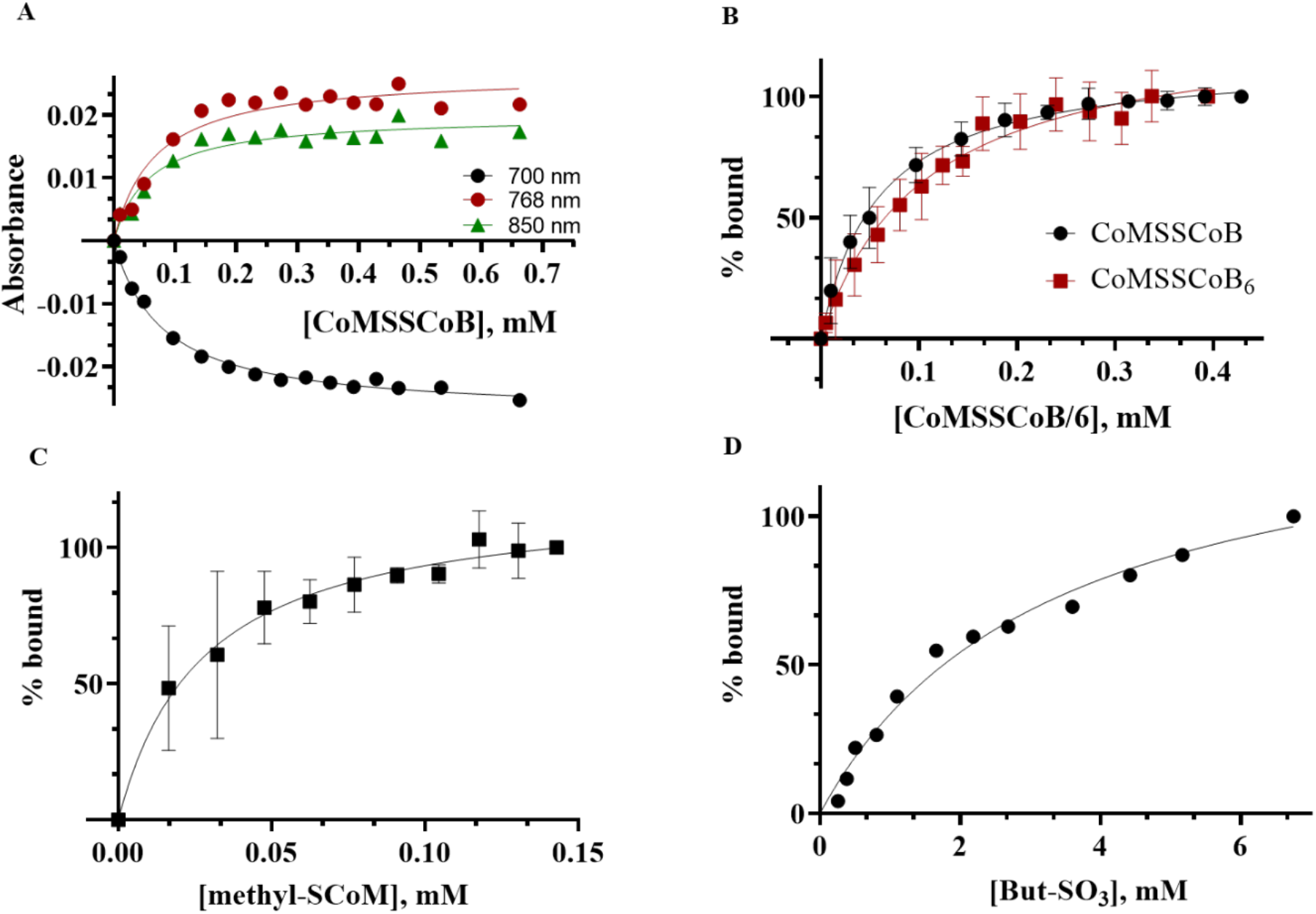
Determination of dissociation constants for MCR substrates and analogs based on NIR spectral changes. A: Plot of changes in absorbance at 700 nm (black), 768 nm (red), 850 nm (green) with increasing concentrations of CoMSSCoB. B: Plot of percent enzyme bound to substrate with addition of increasing concentrations of CoMSSCoB (black) and CoMSSCoB_6_ (red). The data was fit to a one site specific binding equation to obtain K_d_ = 57.4 ± 5.4 µM and 105.3 ± 10.9 µM for CoMSSCoB and CoMSSCoB_6_ respectively. C and D: One site specific binding isotherm for methyl-SCoM gives a K_d_ = 27.53 ± 7.6 µM and But-SO_3_ gives a K_d_ = 3.3 ± 0.6 mM, respectively.

The only structural feature in common among methyl-SCoM and the heterodisulfide substrates relevant to Ni coordination is their sulfonate group (although the sulfurs are similar—thioether and disulfide), indicating the formation of upper axial Ni-O (sulfonate) ligand with these three substrates. Thus, we studied MCR binding to butane sulfonate, which maintains the carbon chain and the sulfonate group but lacks the –S atom that all canonical mechanism (as shown in Fig. 3a) postulate to bind to the Ni(I) during first step of catalysis of both forward and reverse methane synthesis reactions. Upon addition of butane sulfonate, the same NIR spectral shifts are observed as with the other substrates, but with a much higher K_d_ value of 3.3 ± 0.6 mM (**Figure. 9D**). This high value of K_d_ can be due to the lack of the –S which is stabilized via hydrogen bond interactions with the Tyr-333 residue in the catalytic pocket.

### Measurement of pK_a_ values for the sulfonate, carboxylate, phosphate and thiol groups by acid base titrations and ^31^P- and ^1^H-NMR

As a primary standard, 0.1 M NaOH was added to substrate solutions starting at pH 1. The pH changes were recorded and plotted against volume of added 0.1 M NaOH to obtain titration curves. Then, pK_a_ values were calculated using the Henderson-Hasselbach equation (Eq 2) where the pK_a_ equals the pH at half equivalence point.

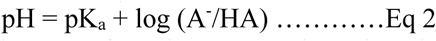

The pK_a_ of 9.4 in CoMSH is assigned to the thiol group. The sulfonate group pKa of 1.9 in CoMSH and 2.5 in methyl-SCoM is as expected due to the inductive effect of the methyl group (electron donating) which destabilizes the conjugate base thus increasing the pK_a_ (**Figure S1**). The pH titration of HSCoB starting at pH 1.1 yields four pK_a_ values (**Table 2A**) that can be attributed to phosphate ionization-1, carboxylate, phosphate ionization-2 and the thiol group (**Figure S2**).

**Table 2.** pKas of ionizable groups of the susbtrates as measured by pH titrations and NMR A.

| pH Titrations - pKas |  |  |  |  |  |
| --- | --- | --- | --- | --- | --- |
| Substrate | -SO<sub>3</sub>H | OPO(OH)(OH) | -COOH | OPO(OH)O- | -SH |
| CoMSH | 1.9 |  |  |  | 9.4 |
| Methyl-SCoM | 2.4 |  |  |  |  |
| CoMSSCoB |  | 1.8 | 4.2 | 7.3 |  |
| CoMSSCoB<sub>6</sub> |  | 2.4 | 4.4 |  | 9.1 |

B.
|  |  | <sup>31</sup>P-NMR | <sup>1</sup>H-NMR |  |  |
| --- | --- | --- | --- | --- | --- |
|  |  | pKa | pKa (g-<sup>1</sup>H) | pKa (h-<sup>1</sup>H) | pKa (k-<sup>1</sup>H) |
| CoMSSCoB | -SO<sub>3</sub>H |  |  |  | <1.25 |
|  | OPO(OH)(OH) | 2.4 |  |  |  |
|  | OPO(OH)O- | 6.6 | 6.2 | 6.8 |  |
|  | -COOH |  | 4.3 |  |  |
| CoBSH | OPO(OH)(OH) | 4.8 |  |  |  |
|  | OPO(OH)O- | 6.6 | 6.4 | 7.2 |  |
|  | -COOH |  | 4.1 |  |  |

To distinguish between the carboxylate and phosphate ionization-1, ^31^P-NMR and ^1^H-NMR chemical shift changes with changes in pH were monitored. The pKa of 9.2 can be unambiguously ssigned to the –SH group of HSCoB (**Figure S3**) (**Table 2B**). In CoMSSCoB, the pKa of the sulfonate group could not be determined because it is presumably lower than 1. The pKa of 1.8 in the case of the heterodisulfide is assigned to the first ionization of phosphate based on the ^31^P-NMR. **Figure S4**

### X-ray absorption studies

XAS studies, including Ni-K edge, pre-edge, and EXAFS of MCRred1 and MCRred1-silent with and without heterodisulfide substrates CoMSSCoB and CoMSSCoB_6_ were carried out to establish the changes in coordination around the Ni(I) in the enzyme substrate binding pocket.

MCRred1 was purified as MCRred1c (CoMSH bound) and was extensively buffer exchanged with 50 mM Tris-HCl, pH 7.6, in an attempt to remove any bound Ni(I)-SCoM and is denoted as MCRred1 in the XAS samples. The Ni(II)-MCRred1-silent sample was prepared by buffer exchange (in 50 mM Tris-HCl, pH 7.6) of an MCRred1c sample, auto oxidized by prolonged storage in the anaerobic chamber. This MCRred1-silent is referred to as MCRred1c-silent going forward in the manuscript for clarity. Based on the UV-visible spectra, the MCred1 XAS spectra were corrected for the presence of 30% MCRred1c-silent impurity present in the sample.

Ni K-edge XAS and EXAFS of MCRred1 show that the Ni(I) is penta-coordinate with 4N (ring) and 1O (axial Gln) (**Figure 10 A and Table 3)**. On the other hand, MCRred1c-silent consists of a hexa-coordinate Ni(II) with 4N (ring), 1O (axial Gln) and a –S ligand from covalently bound CoMSH. The covalent nature of the Ni(II)-S bond prevents the loss of CoMSH during extensive buffer exchange. This is also evident in the EXAFS of MCRred1c-silent where there is a higher contribution from the Ni-S bond and the data can be assigned to a hexa-coordinated Ni(II) (Figure 10A, Table 3)

**Table 3.** Bond distances from XAS.

|  | Ni-N/O Å<br>(penta-coordinate) | Ni-N/O Å<br>(sixth ligand) | Ni-S Å<br>(sixth ligand) from<br>MCR red1 silent |
| --- | --- | --- | --- |
| MCRred1 | 2.03 (4) N ring<br>(1) O Gln |  | 2.39 (0.3) –SHCoM |
| MCRred1 +<br>CoMS-SCoB | 2.04 (4) N ring<br>(1) O Gln | 2.16 (0.7) -O-SO<sub>2</sub>R | 2.39 (0.3) –SHCoM |
| MCRred1 +<br>CoMS-SCoB<sub>6</sub> | 2.03 (4) N ring<br>(1) O Gln | 2.16 (0.7) -O-SO<sub>2</sub>R | 2.39 (0.3) –SHCoM |
| MCRred1c-silent | 2.08 (4) N ring<br>2.20 (1) O Gln |  | 2.42 (1) –SHCoM |
| MCRred1 +<br>CoMSH +HSCoB<br>(partial formation of<br>MCRred2) | 2.05 (4) N ring<br>(1) O Gln | 2.18 (0.3) | 2.45 (0.7) * |

\* XAS is unable to distinguish between MCRred1c-silent and MCRred2, Ni-S interaction. All MCRred1 data were corrected for the presence of a 30% MCRred1c-silent as a contaminant.

**Figure 10.**
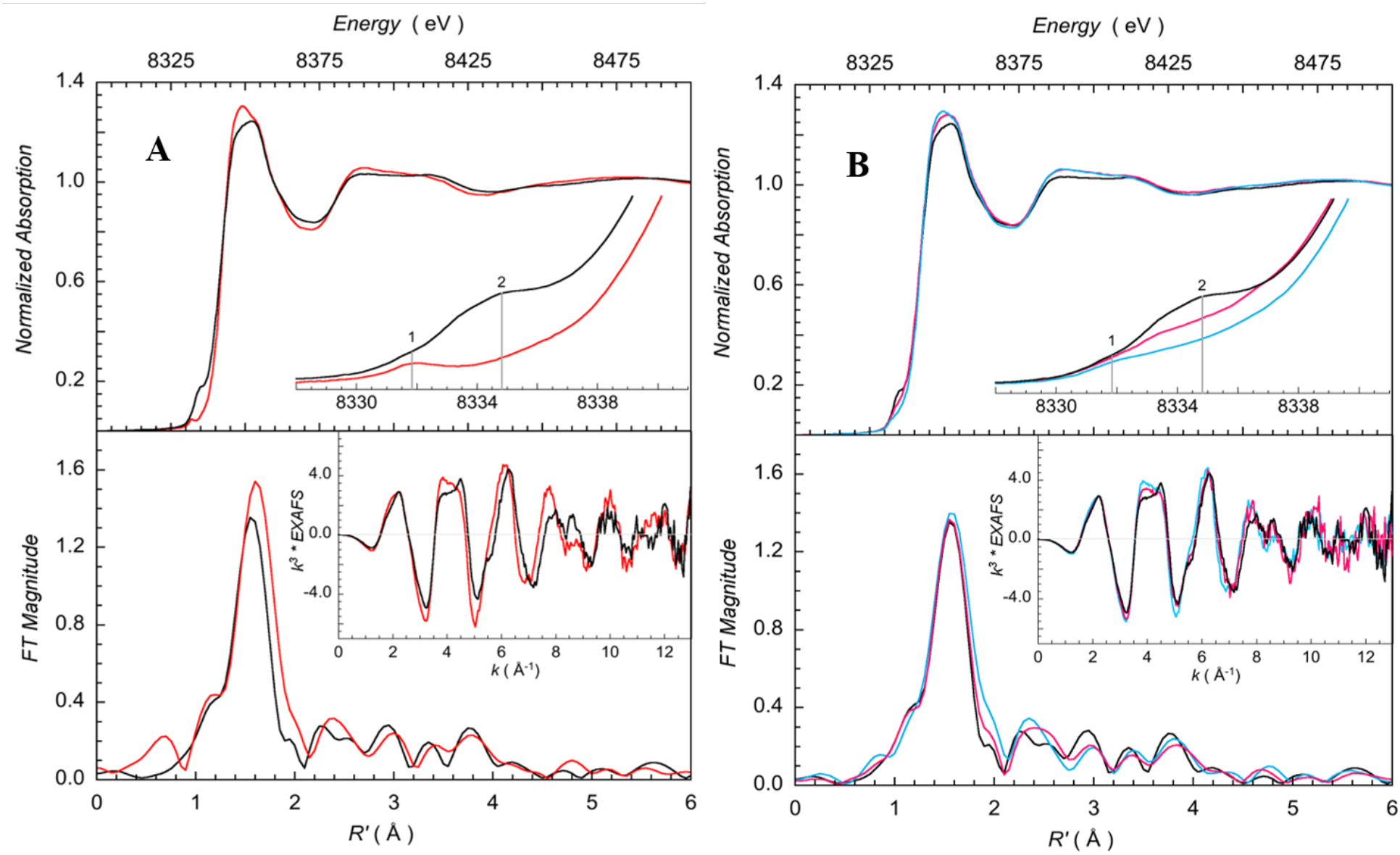
Ni K-edge XAS of MCR + CoBSSCoM. Top row: A comparison of the normalized Ni K-edge XAS data for (A) MCR_Red1_ (—) and MCR_Red1Silent_ (—) and (B) MCR_Red1_ (—), MCR_Red1_ + CoMSSCoB (—) and MCR_Red1_ + CoMSSCoB6 (—). The inset in each plot shows the expanded pre-edge region. The markers at ‘∼8332 eV’ and ‘∼8334.5 eV’ represent the 1s → 3d transition and the back-bonding transition involving interactions between the F430 ring and the Ni center, respectively. Bottom row: A comparison of the Ni K-edge EXAFS data (inset) and their corresponding Fourier Transforms.

Addition of substrates, CoMSSCoB and CoMSSCoB_6_ to MCRred1 results in: a) decrease in the back-bonding contribution at ∼8334.5 eV in the Ni-pre edge region (**Figure 10 B**) and a modest increase in the EXAFS and FT data. FEFF fits suggest the presence of a sixth, weaker, light atom (O/N) ligand to the Ni(I) at ∼2.17 Å (**Figure 11 A and 11 B top panel**). Attempts to fit the data without the weak Ni-O component (**Figure 11A and 11 B, bottom panel**) or by increasing the Ni-S coordination led to statistically poorer fits. Thus, the EXAFS data strongly indicate the formation of a hexa coordinated Ni(I) model for MCRred1 plus CoMSSCoB or CoMSSCoB_6_. Both the heterodisulfide substrates are tethered to the top of the substrate binding channel via phosphate and carboxylate groups with the methylene chain of CoB moiety making its way into the channel, as described above and shown in Fig 1. The sulfonate group of the CoM part of the molecule is within bonding distance of the Ni(I) and is likely the sixth ligand. In contrast, addition of the heterodisulfide substrates to MCRred1c-silent shows no changes in the pre-edge or EXAFS of the enzyme (Figure 6), indicating that the heterodisulfide substrate cannot outcompete covalently bound CoMSH in the Ni(II) active site (**Figure S6**).

**Figure 11.**
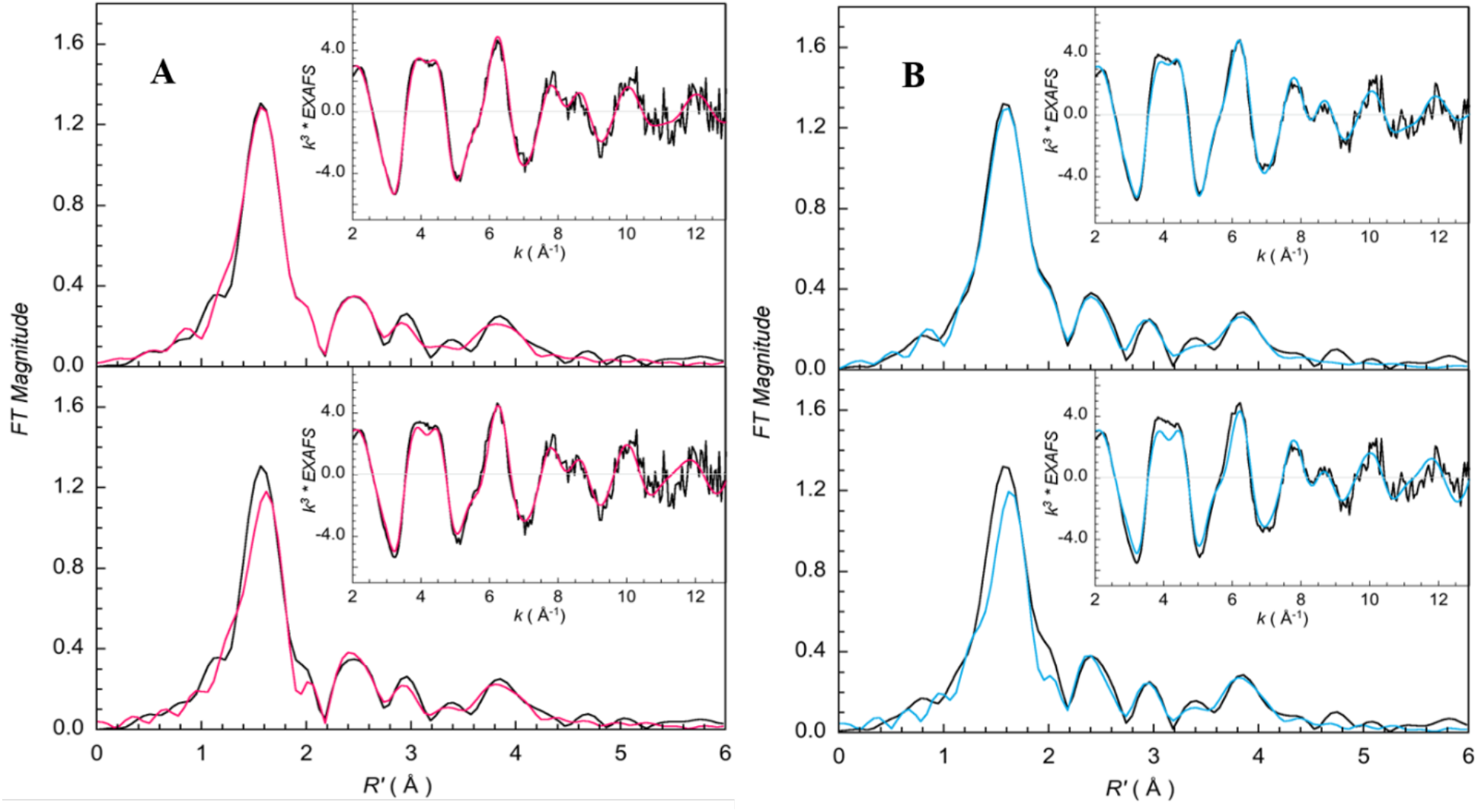
FEFF best fits Ni K-edge EXAFS data (inset) and corresponding Fourier Transforms. The (top) and (bottom) figures represent, respectively, fits with and without a longer Ni-O component in the first shell coordination sphere (see Table 3). A. MCRred1 + CoMSSCoB_6_. Data (—), Fit (—). B. MCRred1 + CoMSSCoB Data (—), Fit (—)

The fits for MCRred1 + CoMSSCoB and CoMSSCoB_6_ are significantly better and statistically relevant when a sixth lighter N/O ligand is included. A Ni(I)-OSO_3_ bond is expected to result in a weak interaction, but the heterodisulfides are constrained in a fixed geometry in the substrate binding pocket with little flexibility. The sulfonate is held in position within binding distance to the Ni(I).

Absence of an increase in the Ni-S component in the FEFF fits to the EXAFS data clearly shows no additional sulfur atom ligation to Ni(I) on addition of any of the substrates. This is quite significant since all catalytic mechanisms begin with a Ni(I)-thiolate. So far, the only experimentally observed Ni(I)-S interaction is with the MCRred2 form, which was generated by addition of HSCoB to MCRred1c. This interaction was previously characterized with pulse EPR and ENDOR studies (35). Similarly, our EPR spectra of the MCRred2 sample (**Figure 12A, bottom spectrum**) shows 40% conversion of MCRred1 to MCRred2 calculated from the relative intensity of the g = 2.2098 (rhombic signal from MCRred2) and g = 2.0537 (axial signal from MCRred1) (Table 3).

EPR spectra of the split XAS samples run in parallel show sharpening of the hyperfine signals arising from the ring nitrogens on addition of CoMSSCoB, CoMSSCoB_6_, methyl-SCoM and But-SO_3_ (**Figure 12 B**) but not with CoMSH (**Figure 12 A bottom spectrum**). The g values (2.2201 and 2.0510) are consistent with previously published EPR data, as summarized (30). As shown earlier (36), methyl-SCoM binding results in sharpening of the superhyperfine features. This sharpening is also observed with addition of CoMSSCoB, CoMSSCoB_6_ and But-SO_3_ suggesting that they bind similarly to MCR. To explain this sharpening of the superhyperfine lines, one option is that it results from movement of the Ni(I) into the plane of the F_430_ corphinoid ring due to the conversion of a penta to a hexacoordinate Ni(I) complex. However, the hyperfine splitting values (*A*_*N*_ and *A*_*Ni*_) of MCRred1 and the methyl-SCoM bound MCRred1m are comparable (37). Perhaps sharpening of the superhyperfine lines results from a Ni(I)-substrate interaction that reduces the micro heterogeneity around the Ni(I) by restricting it to a fixed conformation as the pentacoordinated Ni(I) converts to a hexacoordinated Ni(I) complex with a weak Ni(I)-O-sulfonate interaction. Binding of CoMSH (Figure 12 A, middle spectrum) does not exhibit line sharpening observed with the other substrates, consistent with it not forming an axial Ni(I)-O sulfonate complex.

**Figure 12.**
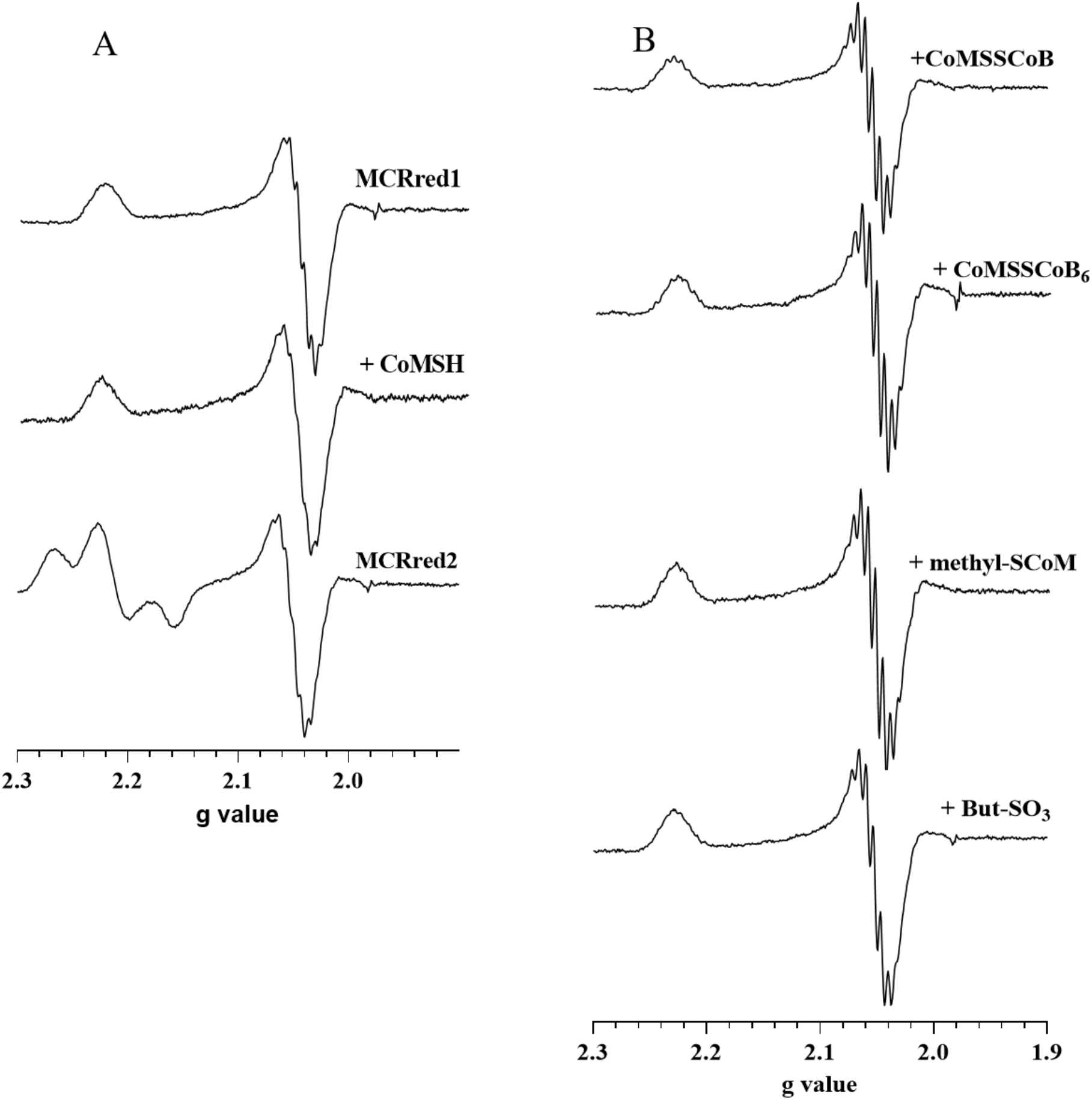
EPR spectra of MCRred1 with substrates. A. MCRred1, MCRred1+ CoMSH, MCRred2 (MCRred1 + COMSH + HSCoB). B. (top to bottom) MCRred1 + CoMSSCoB; CoMSSCoB6; methyl-SCoM; But-SO_3_. Typical EPR signals at 2.2201 and 2.0510 were seen. As shown earlier (36), the superhyperfine features undergo sharpening in the presence of methyl-SCoM but not with CoMSH. The sharpening also is observed with addition of CoMSSCoB, CoMSSCoB_6_ and But-SO_3_ suggesting that they bind similar to methyl-SCoM binding to MCR. However, CoMSH binds differently.

The structural assignments are also consistent with the geometries determined by quantum mechanical calculations of the model systems. The summary of the relevant geometric distances for the three models and the crystal structure 1HBM are summarized in Table 1.

As shown in Fig. 5 and Table 1, the Ni-O distance, between F_430_ and O-Gln147 is similar in the 3 calculated structures at about 2Å, whereas the Ni-O between F_430_ and the SO_3_^-^ group varies, with distance of ∼2.4 for the models 1 and 2, and is elongated to 2.7Å in model 3. Similarly, the distances from the Ni to the S atom in the SO_3_^-^ group is ∼0.3 Å longer in model 3, compared to models 1 and 2. In comparison, the crystal structure shows a longer bond between Ni and Gln147 (2.35 Å), but a slightly shorter distance to the nearest oxygen of the SO_3_^-^ group (2.29 Å). The distances between the Ni center and the S_b_ of HSCoB remains largely unchanged in the three models in Fig. 5 and very similar to the crystal structure; however, the Ni-S_m_ distance varies from ∼6.1Å in CoBSH-methyl-SCoM to ∼ 7Å in CoBSSCoM + CH_4_.

Because no crystal structure is available for methyl-SCoM-bound MCRred1, the position of the CH_3_ group in methyl-SCoM cannot be verified. However, overlap of the calculated structures of CoBSH-methyl-SCoM (Figure 5B) show that even in a state where methyl-SCoM would bind to Ni via the SO_3_^-^ group, the CH_3_ group can fit in a cavity between the Tyr333, Phe330 and HSCoB, nestled near the thiol group of HSCoB (Fig. 5C).

Previous theoretical models that employed a truncated methyl disulfide representation of methyl-SCoM (Fig. 6) where the interaction with Ni is considered via the S_m_ atom, show a Ni-S_m_ distance of ∼3.7Å, and a much longer Ni-O(Gln147) distance of ∼3.8 Å (38). Instead, CoMSSCoB (modeled as ethylS-Smethyl) exhibits a Ni-Sm distance of ∼4Å.

Interestingly, the electronic structure of the Ni center in the present calculations differs from the previous models (19,39). In these prior models, a single unpaired electron is localized on the Ni center, as characterized by Mulliken spin analysis resulting in spin ∼0.9. The lowest electronic state for the present calculations shows that the overall spin on the Ni center is persistently ∼1.6, with an antiferromagnetically coupled electron delocalized on the carbon atoms in the F_430_ ring, giving an overall state Ni(I), even though the metal center itself exhibits Ni(II) like character. The spin assignments for the new structures are reported in **Table 4**. This unusual electronic state of the Ni may explain the uncommon hexa-coordination in the Ni(I).

**Table 4.** Electron spins calculated using Mulliken population analysis at the Ni and select carbons in F_430_ as shown in Figure 5.

| Spin | CoBSSCoM+CH<sub>4</sub> | HSCoB-methyl-SCoM | methyl-SCoM |
| --- | --- | --- | --- |
| Ni | 1.6 | 1.6 | 1.6 |
| C<sup>a</sup> | -0.56 | -0.56 | -0.59 |
| C<sup>b</sup> | -0.39 | -0.39 | -0.35 |

## DISCUSSION

The position of CoBSH in previous crystal structures (29,30) poses a conundrum for all published MCR catalytic mechanisms. Basically, the question is: how can a methyl radical generated from a Ni-thioether interaction with methyl-SCoM at the nickel abstract the H of HSCoB when the reactive sulfurs of the two substrates are 6.4 Å apart? The results described here suggest a solution—methyl-SCoM, the first substrate in the forward reaction, forms a six-coordinate Ni(I)-O complex between a sulfonate oxygen group and the Ni(I) center of F_430_. Similarly, our work suggests that the first substrate in the reverse reaction, CoMSSCoB, forms a Ni(I)-sulfonate interaction.

While the thiolate of CoMS^-^ appears to bind tightly to the Ni(II) states of MCR, in the absence of co-substrate (HSCoB) (Figure 1A), it does not appear to bind to the Ni(I)-state. Furthermore, addition of CoMSH to MCRred1 does not alter its Ni(I) NIR or XAS spectra, indicating that MCRred1 is five-coordinate, with 4N from the hydrocorphin ring and an O from Gln-α’147. This assignment is supported by EXAFS data of the Ni(I)-MCRred1m and MCRred1c states (21). Ni K pre-edge and EXAFS data presented here in addition to time dependent DFT (TD-DFT) calculations also reinforce the five-coordinate nature of the Ni(I) center when MCRred1 reacts with CoMSH (27). Furthermore, when MCRred1 is reacted with ^33^S-substituted CoM^33^SH, no line broadening in the X-band EPR spectrum is observed (28).

Unlike the neutral effect in the NIR spectra of adding CoMSH to Ni(I)-MCR, addition of methyl-SCoM, butylsulfonate, and the heterodisulfides CoMSSCoB and CoMSSCoB_6_ to MCRred1 elicits changes marked in the d-d transitions of the Ni(I) center that reveal the K_d_ values for each of the substrates. Our NIR studies are complemented by XAS, quantum mechanical calculations and TD-DFT studies to characterize the Ni(I) ligand interaction that arises on binding. Furthermore, unlike computational models described so far, which omitted the sulfonate group of methyl-SCoM using methane thiol as the substrate (38), we included this in our computations.

Based on these studies, we propose an alternate model for methyl-SCoM binding to MCRred1 for the forward reaction (**Figure 13**) and a new mechanism (Fig. 3B) for the catalysis of MCR in the forward and reverse reactions. These suggestions depart from canonical mechanisms, e.g. Fig. 3A, which propose a Ni-S (thioether) interaction with methyl-SCoM and a Ni-S (disulfide) interaction with CoMSSCoB. Our alternative model mimics the structure of the CoMSSCoB-product complex with Ni(II)-MCR and positions the methyl thioether group adjacent to the hydrogen atom of HSCoB, as shown in Figure 5.

**Figure 13:**
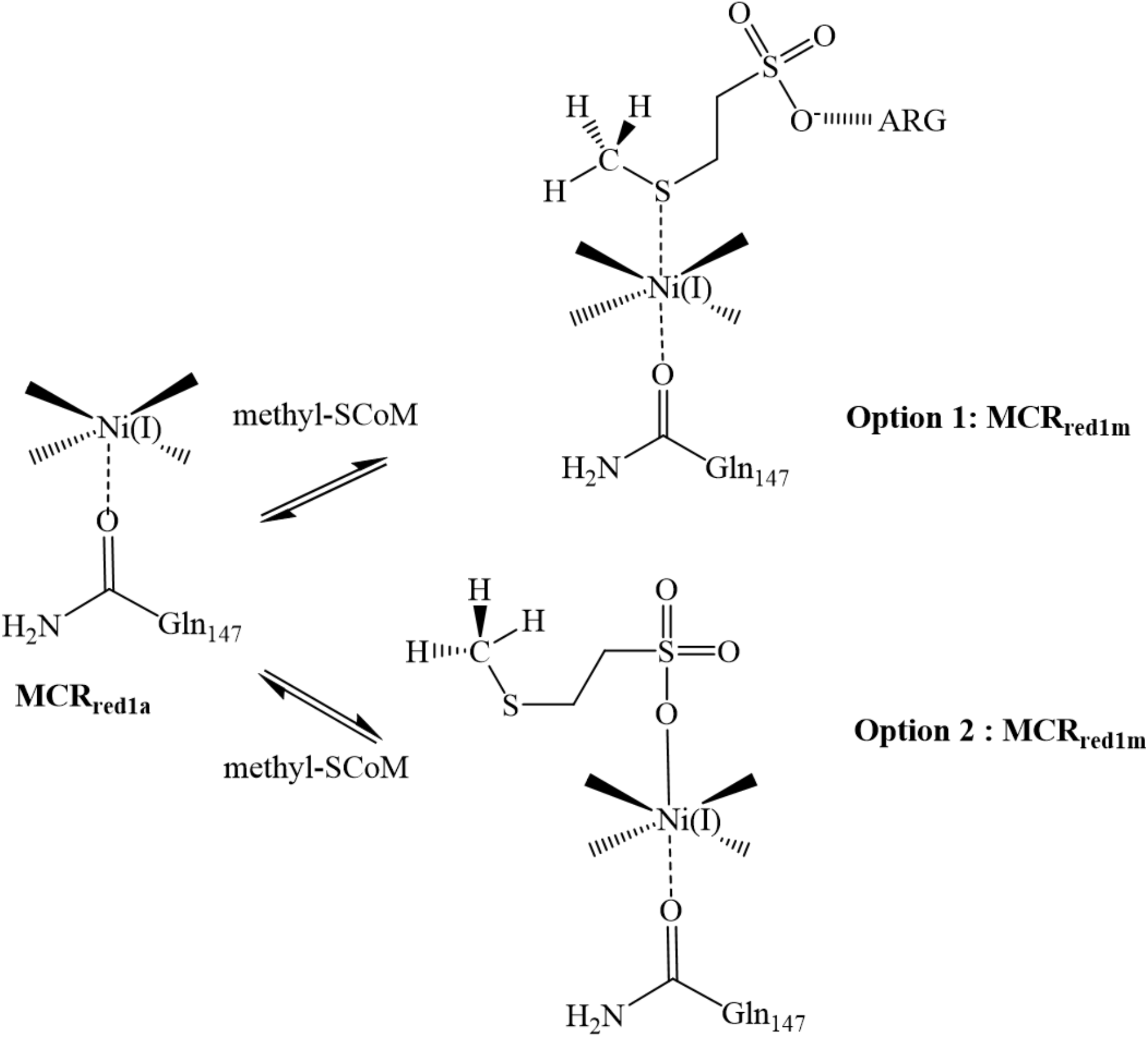
Proposed alternative binding mode for methyl-SCoM to MCR red1. Option 1 highlights the current mode of binding widely accepted as being through the –S of the thioether. Option 2 is our proposed binding mode as seen by NIR spectroscopy for binding of a sulfonate -O^-^ to Ni(I).

Thus, as shown in Figure 5, we propose that the structures of the Ni(I)-methyl-SCoM/HSCoB and the CoMSSCoB complexes mimic that of the X-ray crystal structure of the MCR-silent state, in which the heterodisulfide binds with one –O of the sulfonate axially coordinated to the Ni(II) center and makes H-bond contacts with Tyr-α333. The second –O is hydrogen bonded to the lactam ring of the cofactor F_430_ and to the hydroxyl group of Tyr-β367 and the third –O interacts with a water molecule. Similarly, we propose that the CoBS-part of the heterodisulfide substrate makes the same electrostatic and Van der Waals contacts as the HSCoB at the top of the channel and along the annular arrangement of the hydrophobic residues along the substrate channel. The –S of the CoB-moiety of the heterodisulfide is then placed in the same position as that of HSCoB in the crystal structure. (20)

Thus, in **Step 1** of both the canonical (Fig. 3A) and our proposed (Fig. 3B) mechanisms, Ni(I)-MCRred1 (**A**) first binds methyl-CoM and then CoBSH to form the productive ternary complex **(B)** (34). The shaded box(es) in 3A will be addressed below.

In **Step 2**, Ni(I) transfers an electron to methyl-SCoM to promote homolytic cleavage of the C-S bond of methyl-SCoM forming intermediate (**C)** with the H atom of HSCoB positioned near the CH_3_ group of methyl-SCoM. In Fig. 3A, the proposed transition state (TS1) for this reaction (38) features a Ni-S (thioether) interaction; however, as described above, prior computational models omitted the sulfonate group of methyl-SCoM, thus, its involvement in the MCR mechanism obviously could not be addressed. In Fig. 3B, the methyl-SCoM is flipped 90 degrees with Ni(I)-O (sulfonate) coordination. In both cases, a methyl radical is positioned between the thiolate sulfurs of CoMS- and the H of HSCoB. The mechanistic implication is in the mode of ET—direct Ni→thiolate in 3A and long distance from the sulfonate to the thioether in 3B.

While there has been active discussion in the biology, chemistry and physics communities about how it occurs, long-range ET (aka electron tunneling) of 20 Å has been demonstrated in many proteins (40), such as ribonucleotide reductase (41,42). Redox equivalents can be transferred even longer distances by multistep tunneling, often called hopping, through intervening amino acid side chains. The well-studied mechanism of proton coupled ET (called long-range radical transfer) in ribonucleotide reductase (42) is also instructive.

In **Step 3** of methane synthesis, the methyl group abstracts a hydrogen atom from HSCoB to generate methane and a CoBS• radical (**Species D**). So far we have observed only small amounts (∼6%) of a radical that we have assigned to CoBS• (43) presumably because unstabilized thiyl radicals are difficult to observe due to their short lifetimes and large spin-orbit coupling with the sulfur atom (44). Frey and others have used radical clocks and substrate analogs that form stabilized radicals to investigate elusive substrate radicals (45). The proposed methyl radical, which will likely be even more challenging to visualize, also has not yet been spectroscopically observed.

In **Step 4** of methane synthesis, the CoBS• radical reacts with bound CoM (through the thiolate, 3A, or sulfonate, 3B) to generate a disulfide anion radical (**Species E**), which transfers an electron back to Ni(II) to generate the Ni(I)-CoMSSCoB product (**F**), which we have observed by NIR and XAS spectroscopies. Dissociation of the heterodisulfide in step 5 regenerates the Ni(I) starting state **(A**) for the next round of catalysis.

To address the shaded portion of Fig. 3A, while Fig. 3B invokes the Ni-sulfonate as the productive complex, an alternative is that the Ni(I)-sulfonate complex undergoes rearrangement to a productive Ni-S(thioether) coordination, as proposed in the canonical mechanism. We feel this rearrangement is unlikely because conversion from Species **A** to **B** is quantitative and because our results (described above) indicate that Species **B** (Fig. 3B) is catalytically competent.

## METHODS

### Organism and materials

*Methanothermobacter marburgensis* was obtained from the Oregon Collection of Methanogens (Portland, OR) catalog as OCM82.

All buffers, media ingredients, and other reagents were acquired from Sigma. The N_2_ (99.98%), CO (99.99%), H_2_/CO_2_ (80%/20%), and Ultra High Purity (UHP) H_2_ (99.999%) gases were obtained from Cryogenic Gases (Grand Rapids, MI).

A stock solution of 83 mM Ti(III) citrate was prepared by adding 0.5 M sodium citrate to Ti(III) trichloride (15% w/v in 2N hydrochloric acid) under anaerobic conditions and adjusting the pH to 7.0 with 1M Tris pH 8.0 (46). The concentration of Ti(III) citrate was determined from its UV-visible absorbance at 340 nm (ε= 730 M^-1^ cm^-1^).

Methyl-SCoM was prepared from HSCoM and methyl iodide (47). The homodisulfides CoBS-SCoB and CoB_6_SSCoB_6_ were synthesized from their respective 7-bromoheptanoic acid and 6-bromohexanoic acid (48,49) with a change in the final purification step. RP-PoraPak column was used with an AKTA PURE FPLC system to purify the final homodisulfide with a water/methanol gradient. The heterodisulfide CoMSSCoB7/6 was synthesized via disulfide exchange between HSCoM and CoBSSCoB or CoB_6_SSCoB_6_ in anaerobic 400 mM potassium phosphate buffer for 2h at 45°C. The reaction was stopped by exposure to oxygen followed by purification on a RP-PoraPak column connected to an AKTA PURE FPLC system. The free thiol form of HSCoB and HSCoB_6_ was generated by the reduction of the homodisulfide with tris(2-carboxyethyl) phosphine TCEP in an anaerobic chamber, purified using a RP-PoraPak column and confirmed by HPLC (Phenomenex RP-Fusion C18 column) with a 10mM potassium phosphate (Buffer A) and Acetonitrile (Buffer B) gradient and Mass Spectrometry

### Cell growth and purification

Solutions were prepared and all steps of purification were performed under anaerobic conditions in a Vacuum Atmospheres (Hawthorne, CA) anaerobic chamber maintained under nitrogen gas at an oxygen level below 1 ppm. *M. marburgensis* was cultured on H_2_/CO_2_ (80/20%) at 65 °C in a 14-liter fermenter (New Brunswick Scientific Co., Inc., New Brunswick, NJ) to an optical density of 7-8 at 600 nm in order to increase the yield of MCR-isoenzyme I vs MCR-isoenzyme II. Culture media were prepared as previously described (50) with a slight modification of the sulfur and reducing source, by adding 50 mM sodium sulfide (instead of H_2_S gas) at a flow rate of 1 ml/min during the entire growth period. The cells were anaerobically harvested, resuspended in 50 mM Tris-HCl, pH 7.6, containing 10 mM HSCoM and 0.1 mM Ti(III) citrate, and transferred into a 1-liter serum-stopped anaerobic high-pressure bottle. The headspace of the bottle containing the resuspended cells was purged with CO for 4 hours at 30°C to generate the active MCRred1state as previously described (51). MCR-I was purified from MCR-II using Q-Sepharose Fast Flow resin packed in a XK 16/20 column compatible with AKTA PURE FPLC system. The buffer used was 50 mM Tris-HCl, 10 mM HSCoM pH 7.6 (Buffer A) and 50 mM Tris-HCl, 10 mM HSCoM, 1M NaCl pH 7.6 (Buffer B). Pure MCRred1 was collected in fractions at a gradient centered around 55% Buffer B. The concentration of MCRred1 was determined by UV-visible spectroscopy using extinction coefficients of 27.0 and 9.15 mM^-1^cm^-1^ at 385 and 420 nm, respectively, using a multiple wavelength calculation as previously described. (50) The concentration of MCRred1-silent, which contains the inactive Ni(II) form of F_430_, was calculated using extinction coefficients of 22.0 and 12.7 mM^-1^cm^-1^at 420 and 385 nm, respectively (50). This purification method yields between 65-75% MCRred1 in the active Ni(I) form and is used as is in all experiments unless otherwise stated.

### UV-visible, NIR and EPR studies

Absorbance spectra were recorded in the anaerobic chamber using a diode array spectrophotometer HP-8453 instrument. EPR spectra were recorded on a Bruker EMX spectrometer (Bruker Biospin Corp., Billerica, MA), equipped with an Oxford ITC4 temperature controller, a Hewlett-Packard model 5340 automatic frequency counter, and Bruker gauss meter. The EPR spectroscopic parameters included the following: temperature, 100K; microwave power, 10 milliwatt; microwave frequency, 9.43GHz; receiver gain, 2104; modulation amplitude, 10.0 G; modulation frequency, 100 kHz. Spin concentration was determined by double integration of the sample spectrum obtained under non saturating conditions and comparison to that of 1 mM copper perchlorate standard. All samples for EPR spectroscopy were prepared in 50 mM Tris-HCl, pH 7.6, in a Vacuum Atmospheres anaerobic chamber.

### Determination of dissociation constants

The interaction of substrates with MCR was determined by monitoring changes in the Near-IR spectrum of active Ni(I) in MCR red1. The enzyme used was prepared by removing HSCoM and Ti(III) citrate from MCR by buffer exchange with 50 mM Tris-HCl, pH 7.6, using Amicon Ultra15 centrifuge filter units with a 30-kDa cut-off (Millipore).

CoMSSCoB, CoMSSCoB_6_, or methyl-SCoM (0-500 µM) was added in small increments to 50 µM MCR_red1_ and the changes in the NIR spectra were measured. (0-10 mM) But-SO_3_ or HSCoM was added to 50 µM MCRred1-Ni(I) in order to monitor the changes induced due to substrate binding. The changes in absorbance (700 nm, 768 nm and 850 nm) were plotted against substrate concentration and fit to a one site binding isotherm to obtain the dissociation constants for the various substrates. Instantaneous changes in absorbance were recorded in order to study binding and the titration data was collected over 5 min wherein no redox change with Ni(I) was observed.

### XAS

The Ni K-edge XAS studies on MCR were measured at the Stanford Synchrotron Radiation Lightsource (SSRL) on the unfocused 20-pole 2 T wiggler sidestation beamline 7-3 under nonstandard ring conditions of 3 GeV and ∼500 mA (low-alpha operations mode at SSRL). A Si(220) double crystal monochromator was used for energy selection. The no-M0 mirror configuration was used and components of higher harmonics were rejected by detuning the monochromator by ∼30%. All samples were measured as solutions, which were transferred to 1 mm Delrin XAS cells with 1 mil Kapton tape windows under anaerobic conditions and were immediately frozen after preparation and stored under liquid N_2_. During data collection, samples were maintained at a constant temperature of ∼10 K using a closed-cycle CryoIndustries liquid Helium cryocooler. Data were measured to k = 13 Å^-1^ (fluorescence mode) using a Canberra Ge 30-element array detector. Internal energy calibration was accomplished by simultaneous measurement of the absorption of a Ni foil placed between two ionization chambers situated after the sample. The first inflection point of the foil spectrum was fixed at 8331.6 eV. The samples were monitored for photoreduction and a fresh spot was chosen for data collection after every three scans. However, no visual change in the rising edge energy position or the pre-edge features (associated with the Ni(I) and Ni(II) forms) were observed over successive scans, indicating that all the samples were resistant to photoreduction under the reduced current experimental conditions. Data presented here have been averaged over 12-scans or more, depending on the species. Data were processed by fitting a second order polynomial to the pre-edge region and subtracting this from the entire spectrum as background. A four-region spline of orders 2, 3, 3 and 3 was used to model the smoothly decaying post-edge region. The data were normalized by subtracting the cubic spline and assigning the edge jump to 1.0 at 8335 eV using the Pyspline (52) program. Data were then renormalized in Kaleidagraph for comparison and quantification purposes.

Theoretical EXAFS signals χ(k) were calculated by using FEFF (Macintosh version 8.4) (53-55). Initial structural model for MCR was obtained from the crystal structure, and modified in Avogadro to consider O- and S-axial ligands (56). The input structure was improved based on preliminary EXAFS fit parameters to generate more accurate theoretical EXAFS signals. Data fitting was performed in EXAFSPAK (57). The structural parameters varied during the fitting process were the bond distance (R) and the bond variance σ^2^, which is related to the Debye-Waller factor resulting from thermal motion, and static disorder of the absorbing and scattering atoms. The nonstructural parameter ΔE_0_ (E_0_ is the energy at which k = 0) was also allowed to vary but was restricted to a common value for every component in a given fit. Coordination numbers were systematically varied in the course of the fit but were fixed within a given fit. The fits to the MCRred1-silent yielded metrical parameters nearly identical to those reported earlier. The presence of a MCRred1-silent in the reduced MCRred1 was accounted for by including a fixed 0.4 coordination number Ni-S path fixed at the distance obtained from the MCRred1-silent fit.

140 µL samples of MCR red1 [70% Ni(I)] 600 µM incubated with various substrates CoMSSCoB (20 mM), CoMSSCoB6 (30 mM), methyl-ScoM (10 mM), But-SO_3_ (40 mM) or HSCoM (40 mM) and MCR red1 [70% Ni(I)] (600 µM) + HSCoM (10 mM) + CoB7SH/ CoB6SH (20 mM) were prepared in 50 mM Tris-HCl pH 7.6 in 40% Glycerol anaerobically. 90 uL of each sample was frozen in a XAS cell and 50 uL of the same sample added to 100 µL of 50 mM Tris-HCl pH 7.6 was frozen in EPR tubes for measurement.

### pKa measurements

pKa measurements of the various substrates was carried out using acid base titrations with 0.1 M standard sodium hydroxide solution (Sigma) to obtain titration curves. ^31^P-NMR and ^1^H-NMR were measured at various pH values for all the substrates and the changes in chemical shifts with pH were recorded. pKa values for the various functional groups (carboxylate, phosphate and thiol) were obtained by plotting chemical shifts as a function of pH.

### Computational studies

Quantum mechanical calculations were done using the NWChem program (58), employing DFT methodology. All structures were optimized using the B3LYP (59,60) functional and the dispersion correction was added using Grimme’s D3 correction (61). All non-metal atoms were represented with the 6-31G* Gaussian basis set (62), and for the Ni center we used lanl2dz ECP basis set (63). The structures were extracted from the crystal structure deposited in the PDB bank under code 1HBM. Points of truncation from the full structure are marked with an asterisks in Figures 5 and 6 where the model is shown. The truncated bonds were terminated with hydrogens. In the geometry optimization procedure, the heavy atoms at the truncated points were fixed in space. The structures were optimized in gas phase.

The UV vis spectra were calculated using TDDFT (64-66) approach in NWChem, using the RPA approximation. The wave functions were optimized at the same level of theory used in the geometry optimization, and a total of 50 excited states were calculated and used in the reconstruction of the spectra, using Gaussian broadening functions.

## ABBREVIATIONS

The abbreviations used are AOM, anaerobic oxidation of methane; HSCoB, Coenzyme B, N-7-mercapto<u>heptan</u>oylthreonine phosphate; HSCoB_6_; N-6-mercapto<u>hexan</u>oylthreonine phosphate; EPR, electron paramagnetic resonance; ENDOR, electron nuclear double resonance; ET, electron transfer; MCD, magnetic circular dichroism; MCR, methyl-CoM Reductase; methyl-SCoM, S-methyl-2-mercaptoethane sulfonate; But-SO_3_, butane sulfonate; CoMSH, 2-mercaptoethane sulfonate; CoMSSCoB, heterodisulfide of coenzyme M and coenzyme B, NIR, near infrared; RR, Resonance Raman spectroscopy; TS, transition state; XAS, X-ray absorption spectroscopy EXAFS, Extended X-ray Absorption Fourier Transform Spectroscopy; NMR, Nuclear Magnetic Resonance Spectroscopy

## ACKNOWLEDGMENTS

The content is solely the responsibility of the authors and does not necessarily represent the official views of the Department of Energy.

## DATA AVAILABILITY

The data are all contained in this manuscript.

