## Supplemental Materials for "Nickel-Sulfonate Mode of Substrate Binding for Forward and Reverse Reactions of Methyl-SCoM Reductase Suggest a Radical Mechanism Involving Long Range Electron Transfer"

Running title: *Substrate binding via nickel-sulfonate to methyl-SCoM reductase*

[1] To whom correspondence should be addressed: Department of Biological, Chemistry, University of Michigan Medical School, 1150 W. Medical Center Dr., 5301 MSRB III, Ann Arbor, MI USA, 48109-0606, Tel: 734-615-4621; Fax: 734-763-4581;

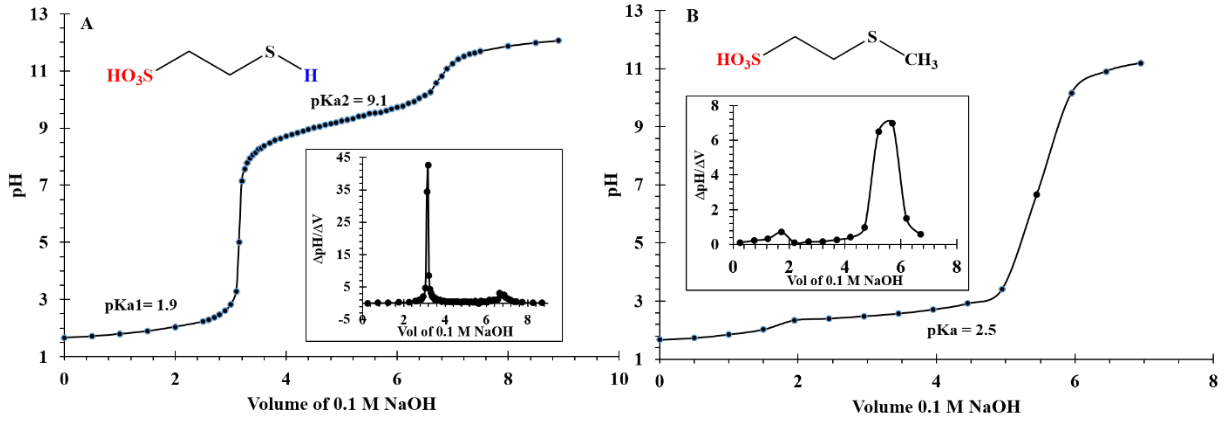


**Figure S1: pH titration curves for MCR substrates.** A: CoMSH and B: methyl-SCoM. Insets correspond to the derivative plot with peak corresponding to equivalence point pH. pH measured at half equivalence point is the pKa.

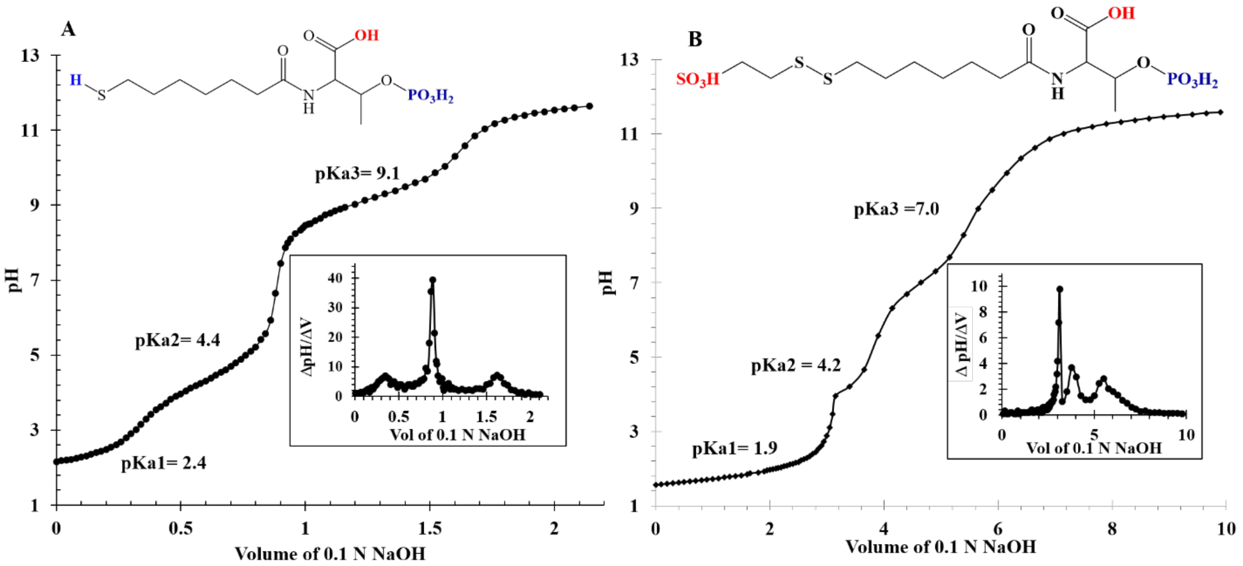


**Figure S2. pH titrations of MCR substrates**. (A) CoBSH and (B) CoMSSCoB. Insets are plots of ratio of change in pH/ change in volume ( ΔpH/ ΔV) vs Vol (V) with peak corresponding to equivalence point volume. pH measured at half equivalence point is the pKa of the group being titrated.

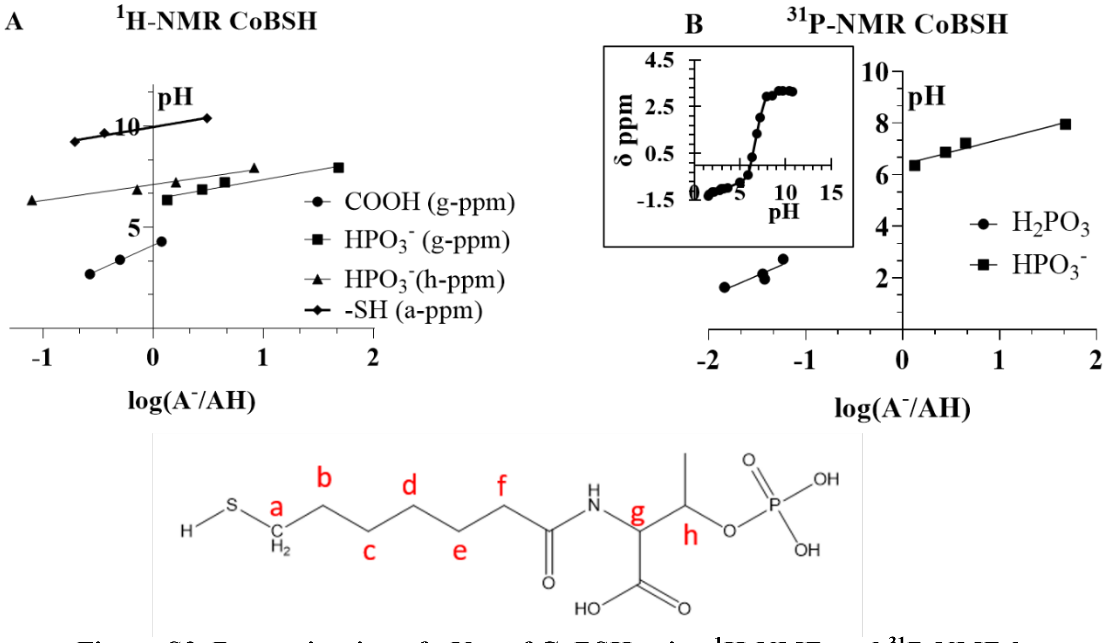


**Figure S3. Determination of pKas of CoBSH using $^{1}$H-NMR and $^{31}$P-NMR by monitoring changes in chemical shifts with pH**. (A) Linear plot of pH vs log ($A^-$/AH) (ratio of deprotonated to protonated species) results in a Y-intercept that gives the pKa of the titrated group. Changes in δ ppm of Hg- proton (sensitive to deprotonation of –COOH), Hh-proton (sensitive to $H_2PO_3$ ionization) and Ha-proton (sensitive to thiol group deprotonation) due to changes in pH are used to determine the pKa of the ionizable group. (B) pKa values of phosphate group of CoBSH as determined by $^{31}$P-NMR measurements. Y intercept for the linear plot of log($A^-$/AH) vs pH is pKa for the corresponding deprotonation. Concentration of $A^-$ and AH is calculated from concentration and volume of 0.1 M NaOH added (67).

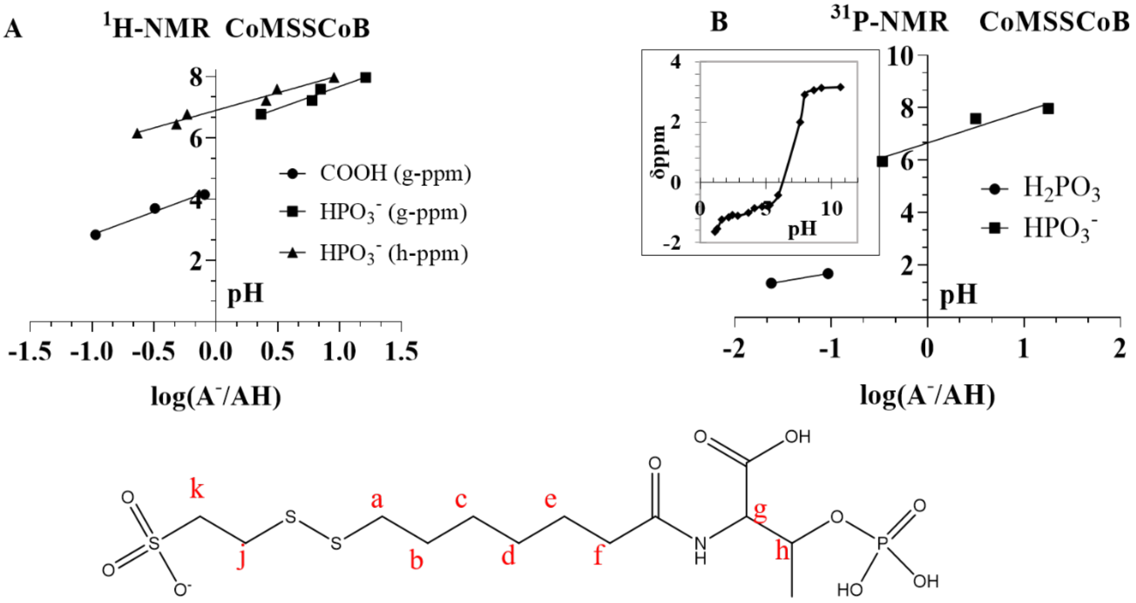


**Figure S4. Determination of pKas of CoMSSCoB using $^{1}$H-NMR and $^{31}$P-NMR by monitoring changes in chemical shifts with pH**. (A) Linear plot of pH vs log ($A^-$/AH) (ratio of deprotonated to protonated species) results in a Y-intercept that gives the pKa of the titrated group. Changes in δ ppm of Hg- proton (sensitive to deprotonation of –COOH), and Hh-proton (sensitive to $H_2PO_3$ ionization) due to changes in pH are used to determine the pKa of the ionizable group. The Hk proton chemical shift remained unchanged throughout the experiment suggesting it was deprotonated at pH 1. (B) pKa values of phosphate group of CoBSH as determined by $^{31}$P-NMR measurements. Y intercept for the linear plot of log($A^-$/AH) vs pH is pKa for the corresponding deprotonation. Concentration of $A^-$ and AH is calculated from concentration and volume of 0.1 M NaOH added.(67)

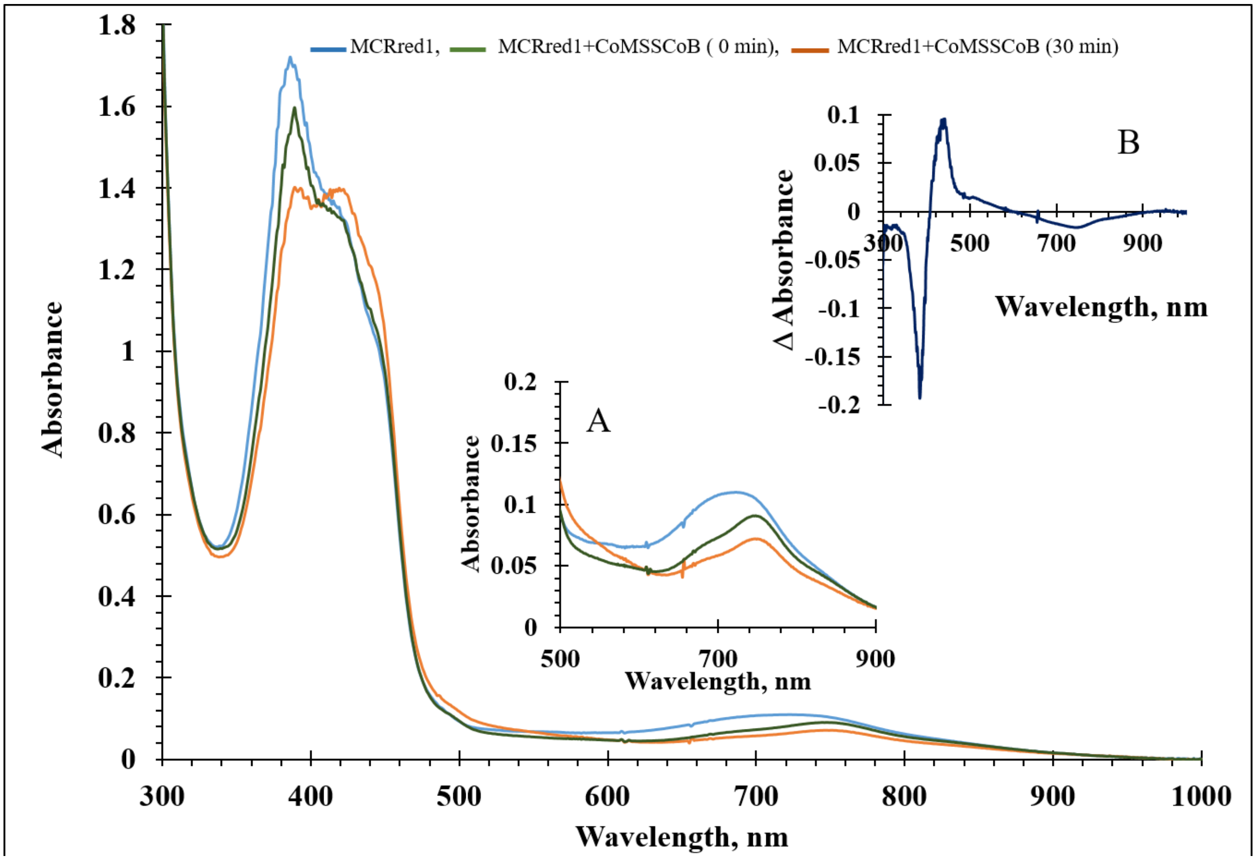


**Figure S5. UV-visible and NIR spectra of reaction of MCRred1 with CoMSSCoB.** Reaction of 50 μM MCRred1 + 5 mM CoMSSCoB monitored from 300nm to 1000nm to capture the changes in redox state of MCRred1 [Ni(I) @ 385 nm → Ni(II)/Ni(III) @ 420/445(sh) nm] over 30 minutes. Inset A. NIR changes highlight the shift in λmax from (—)700nm → (—) 768 nm on addition of CoMSSCoB and its subsequent decay over time (—). Inset B. Difference spectra of the spectra at 30 min and 0 min highlights the increase in absorbance at 445 nm associated with a decrease in absorbance at 768 nm.

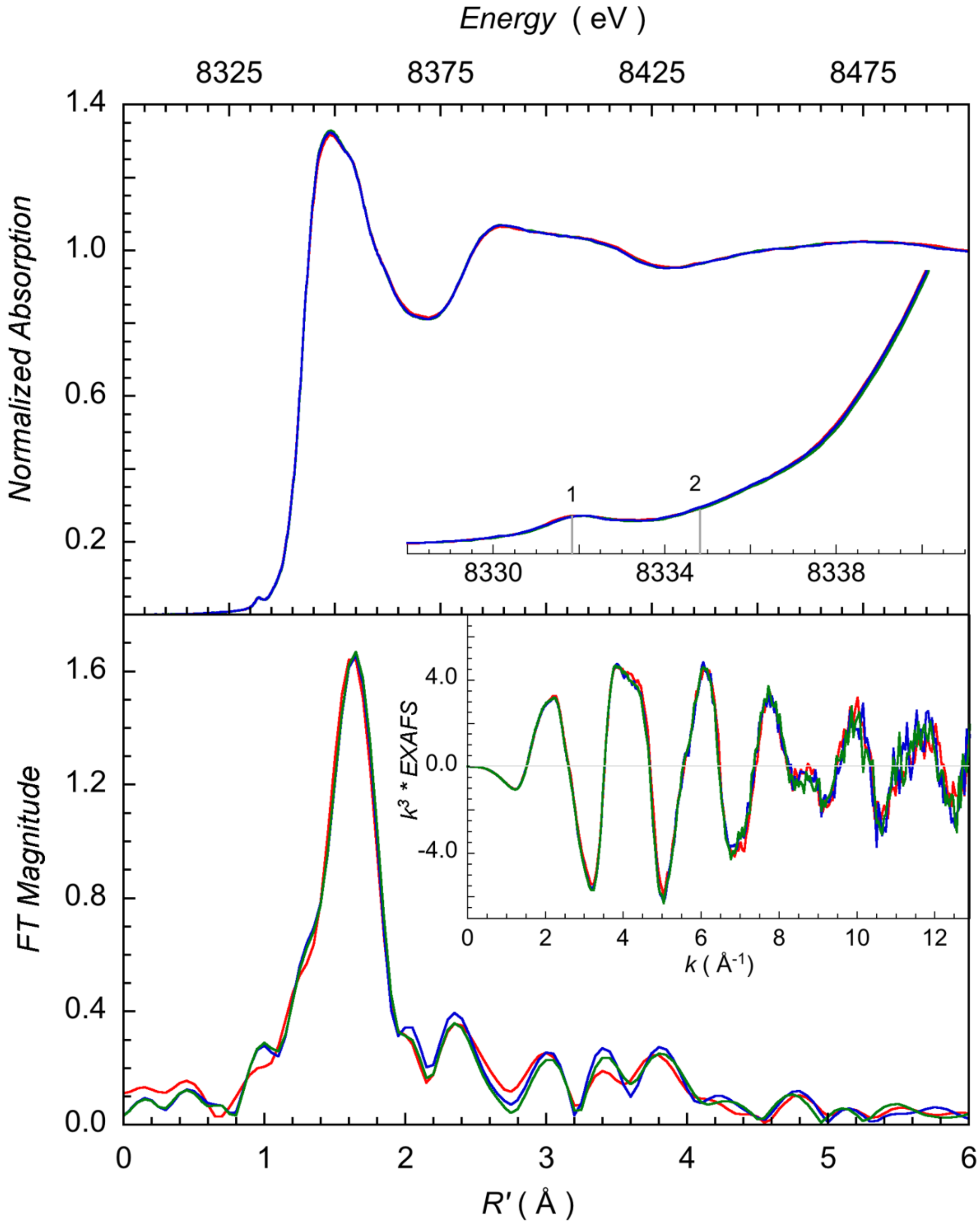


**Figure S6. XAS spectra of MCRred1c-silent with substrates shows no changes.** Top row: A comparison of the normalized Ni K-edge XAS data for $MCR_{Red1Silent}$ (—), $MCR_{Red1Silent}$ + CoMCoB (—) and $MCR_{Red1Silent}$ + CoMCoB6 (—). The inset in each plot shows the expanded pre-edge region. The markers at '~8332 eV' and '~8334.5 eV' represent the 1s $\rightarrow$ 3d transition and the back-bonding transition involving interactions between the F430 ring and the Ni center, respectively. Bottom row: A comparison of the Ni K-edge EXAFS data (inset) and their corresponding Fourier Transforms.
